# Stable Cortical Body Maps Before and After Arm Amputation

**DOI:** 10.1101/2023.12.13.571314

**Authors:** Hunter R. Schone, Roni O. Maimon Mor, Mathew Kollamkulam, Malgorzata A. Szymanska, Craig Gerrand, Alexander Woollard, Norbert V. Kang, Chris I. Baker, Tamar R. Makin

**Affiliations:** Institute of Cognitive Neuroscience, University College London, London, UK; Laboratory of Brain & Cognition, National Institutes of Mental Health, National Institutes of Health, Bethesda, Maryland, USA; Rehab Neural Engineering Labs, University of Pittsburgh, Pittsburgh, PA, USA; Department of Physical Medicine and Rehabilitation, University of Pittsburgh, Pittsburgh, PA, USA; Department of Experimental Psychology, University College London, London, UK; UCL Institute of Ophthalmology, University College London, London, UK; Department of Experimental Psychology, University of Oxford, Oxford, UK; MRC Cognition and Brain Sciences Unit, University of Cambridge, Cambridge, UK; Department of Orthopaedic Oncology, Royal National Orthopaedic Hospital NHS Trust, Stanmore, Middlesex, UK; Plastic Surgery Department, Royal Free Hospital NHS Trust, London, UK; Wellcome Centre for Human Neuroimaging, UCL Institute of Neurology, London, UK

## Abstract

The adult brain’s capacity for cortical reorganization remains debated. Using longitudinal neuroimaging in three adults, followed up to five years before and after arm amputation, we compared cortical activity elicited by movement of the hand (pre-amputation) versus phantom hand (post-amputation) and lips (pre/post-amputation). We observed stable representations of both hand and lips. By directly quantifying activity changes across amputation, we overturn decades of animal and human research, demonstrating amputation does not trigger large-scale cortical reorganization.

---

What happens to the brain’s map of the body when a part of the body is removed? Over the last five decades, this question has captivated neuroscientists and clinicians, driving research into the brain’s capacity to reorganize itself. Primary somatosensory cortex (S1), known for its highly detailed body map, has historically been the definitive region for studying cortical reorganization^1,2^. For example, foundational research in monkeys reported that, following an amputation or deafferentation, the affected region within the S1 body map suddenly responds to inputs from cortically-neighboring body-parts (e.g., face)^3,4^. Additional neuroimaging studies in human amputees supported the theory that amputation of an arm triggers large-scale cortical reorganization of the S1 body map^5–7^, with a dramatic redistribution of cortical resources, hijacking the deprived territory^1^.

Recent studies have challenged this view by harnessing human amputees’ reports of experiencing vivid sensations of the missing (phantom) limb. First, human neuroimaging studies have demonstrated that voluntary movements of phantom fingers engage neural patterns resembling those of able-bodied individuals^8–10^. Second, phantom sensations are evoked by cortical^11^ or peripheral^12,13^ nerve stimulation, suggesting an intact neural representation of the amputated limb, despite its physical absence. Third, neuroimaging studies using both tactile stimulation and movement paradigms reported no changes in face or lip activity within the deprived cortex of adult amputee participants compared to able-bodied controls^14,15^, (though remapping observed in children)^16^.

This debate—whether or not amputation triggers large-scale reorganization— remains unresolved^17,18^, with some suggesting the two views are not conceptually exclusive – preservation and reorganization can co-exist^5,19,20^.

However, a fundamental issue with the evidence on both sides of this debate is a methodological reliance on cross-sectional designs (i.e., comparing between participants). While offering valuable proofs of concept, these studies cannot determine whether the maps of the phantom hand or face are truly preserved or changed, relative to their pre-amputation state. To directly track the evolution of cortical representations before and after amputation, we implemented a longitudinal fMRI approach to track the cortical representations of the hand and face (lips) in three adult participants up to 5 years after arm amputation (Video 1), compared with able-bodied control participants (Figure 1A). Avoiding the confounding effects of cross-sectional designs^21^, we directly quantify the impact of arm amputation on S1 (re)organization.

**Figure 1.**
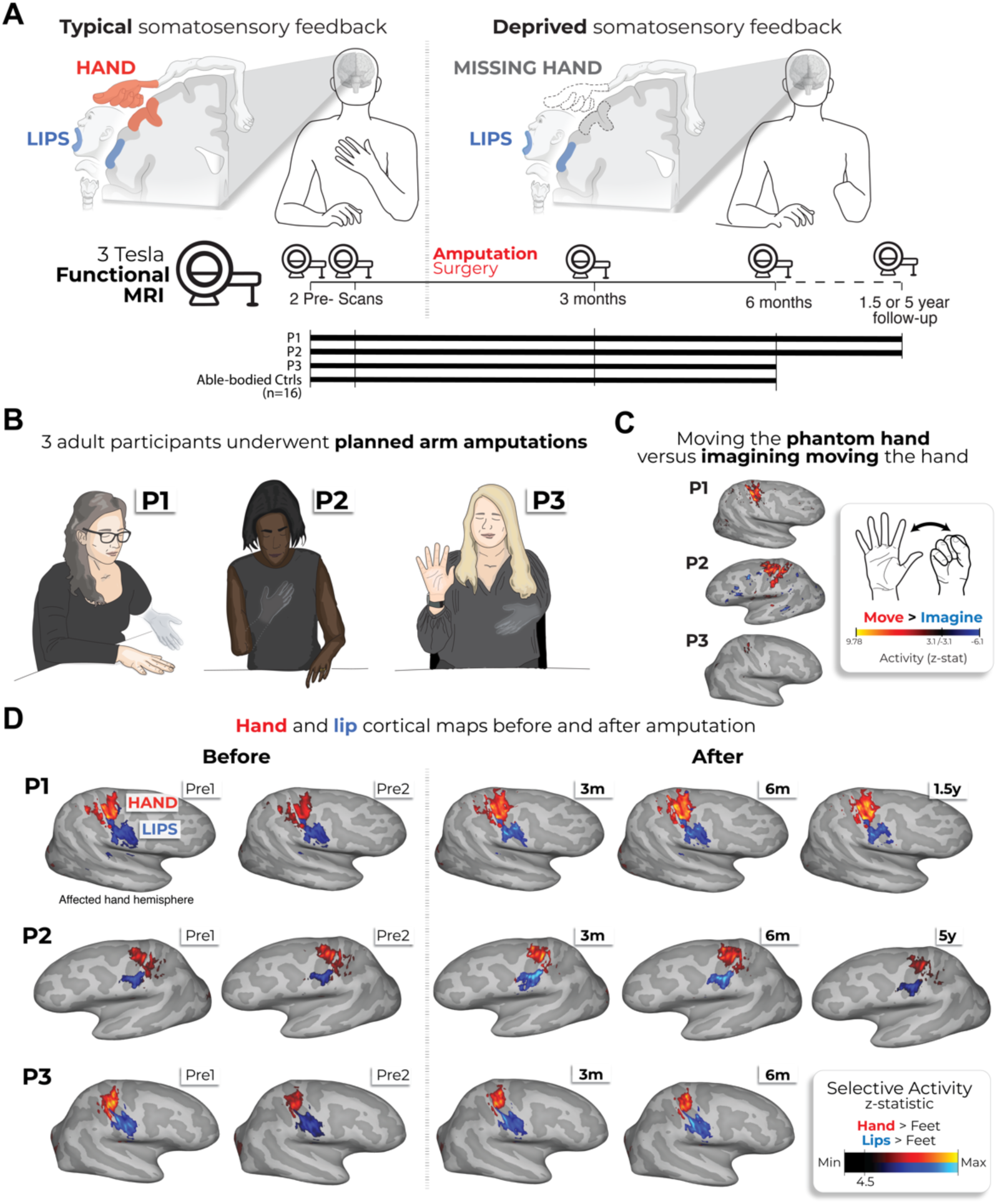
Longitudinal investigation of participants with planned arm amputations. **(A)** Experimental timeline. Pre- and post-amputation scans were conducted across 4-5 time points: twice before, and at 3 months, 6 months and 1.5 (P1) / 5 years (P2) after amputation. **(B)** Illustration depicting the 3 participants 6m post-amputation, including their subjective description of their phantom limb position. **(C)** Phantom movements are not imaginary. Univariate activity (z-scored) contrast map displaying participant’s attempts to open and close the phantom hand vs. imagining movement, 6 months post-amputation. **(D)** Participant’s hand (red) and lip (blue) cortical activation maps (contrasted against feet movements) within the affected hand hemisphere across 4-5 sessions. All maps were minimally thresholded at 33% the maximum z-statistic and used a common color scale (participant’s maximum Z-statistic > 4.5).

We studied three adult participants (case-studies: P1, P2, P3) undergoing arm amputation (demographics in Supp. Table 1) across 4-5 timepoints, and 16 able-bodied controls at 4 timepoints over 6 months (Figure 1A). Pre-amputation, all participants could move all fingers, to varying ranges (Supp. Figure 1 and Supp. Video 1). Post-amputation, all participants reported vivid phantom limb sensations (Figure 1B), including volitional phantom fingers movement (Supp. Table 1 and Supp. Figure 1). Motor control over the phantom hand was further confirmed by residual limb muscle contractions during phantom movements (Supp. Video 1), and selective activation in primary sensorimotor cortex for attempted, but not imagined, phantom movements (Figure 1C). The critical question is to what degree S1 phantom activity reflects the pre-existing hand.

During scanning, participants performed visually-cued movements involving tapping individual fingers, pursing lips, and flexing toes. Case-study participants demonstrated strikingly consistent hand and lip cortical maps before and after amputation (Figure 1D). Projecting hand and individual fingers activity profiles across S1 revealed stable activity before and after amputation, with phantom activity resembling the pre-amputation amplitude and spatial activity spread (Figure 2A). A center of gravity (CoG) analysis of these profiles revealed spatially consistent hand and individual finger activity in our case-studies, with similar pre- to post-session differences over 6 months as controls (six Crawford t-tests per participant: P1: 0.14≤*p_uncorr_*≤0.58; P2: 0.06≤*p_uncorr_* ≤0.81; P3: 0.10≤*p_uncorr_*≤0.91).

**Figure 2.**
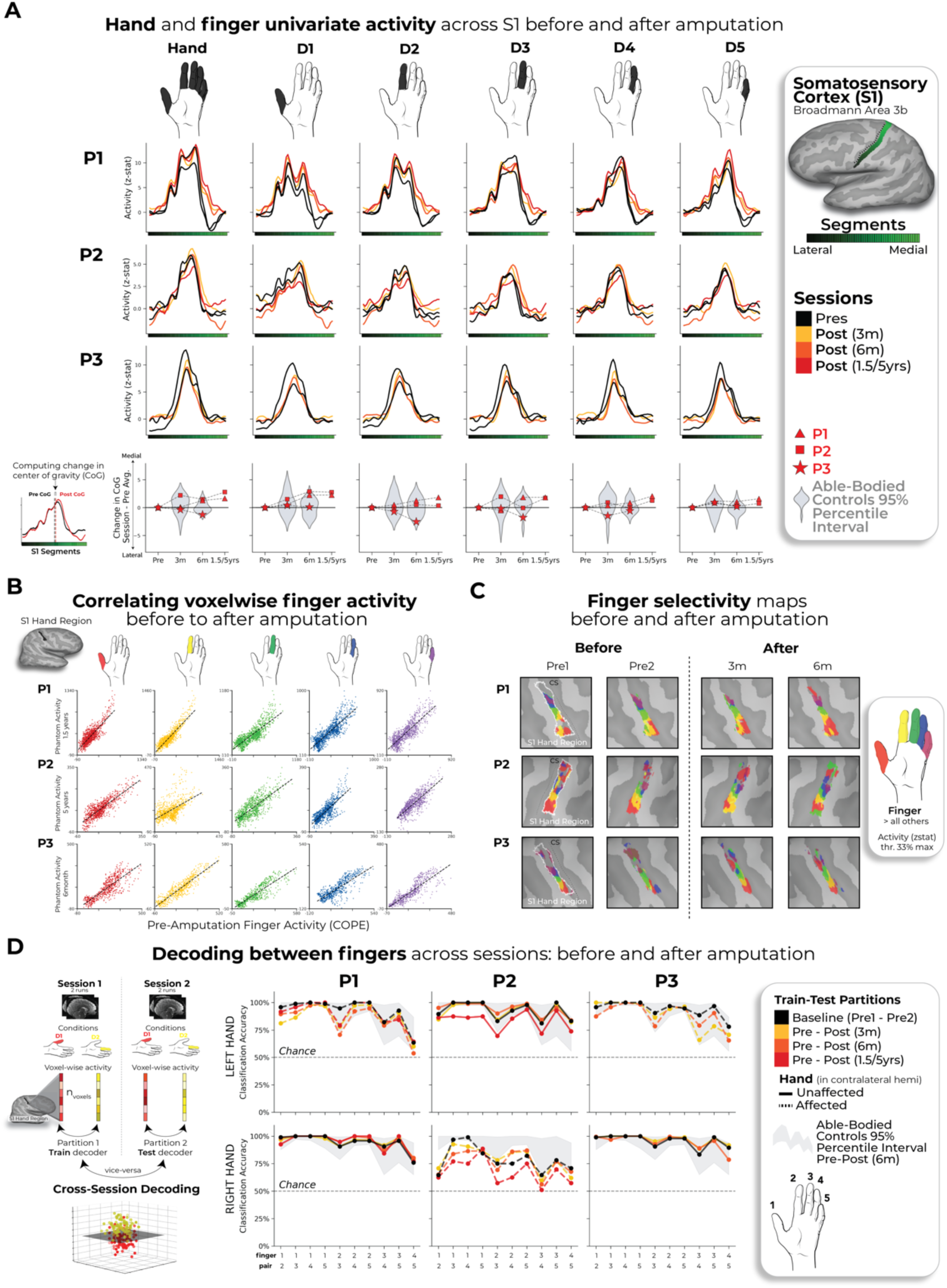
Stable hand representation within the affected hemisphere despite amputation. **(A)** Longitudinal hand and individual finger activity (versus rest) projected across the S1 (BA3b) region of interest (ROI) segmented into 49 segments of similar height. Affected hand’s activity over 5 sessions (indicated in the legend) for each of the case-study participants that underwent an amputation; bottom row shows finger CoG shifts before and after amputation. Black lines reflect pre-amputation activity, orange/red lines post-amputation. Case-study participants’ CoG shifts (red) for the hand and individual fingers fell within the distribution of controls (grey; 12-18 comparisons per participant; Crawford t-tests: P1 (6m): 0.14≤ p_uncorr_≤ 0.58; P2 (6m): 0.06≤p_uncorr_≤0.81; P3 (6m): 0.10≤p_uncorr_≤0.91). Values indicate group means ± standard error. Positive values indicate medial shifts (toward feet), negative values lateral (toward lips) in S1. Control data shown as gray violin plots. P1 data shown as a red triangle. P2 data shown as a red square. P3 data shown as a red star. For simplicity, the control values are all for the left (non-dominant) hand. **(B)** Pre-post amputation single- finger multi-voxel correlation: For each finger of the case-study participants, voxel-wise activity correlations before and at the final scan after amputation are shown. All other correlations are comprehensively reported in Supp Figure 5. All participant’s pre-to-post correlations were significant (5 Pearson correlations per participant; P1 (6m): 0.68≤ r≤ .90, p_uncorr_<0.001; P2 (6m): 0.80≤r≤.85, p_uncorr_<0.001; P3 (6m): 0.88≤r≤.91, p_uncorr_<0.001). **(C)** Finger selectivity maps before and after amputation. Each contrast map reflects the activity for each finger (versus all others), masked to the hand ROI. Each mask was minimally thresholded at 33% the maximum z-statistic. Color codes indicated on the right. To capture the multi-finger activity at a single voxel, a 70% opacity filter was applied to all fingers. **(D) Left** - Graphic illustration of multivoxel analyses using a linear SVM decoder. **Right** – Longitudinal classifier performance. Line colors denote train-test/cross validation session pairs, respectively as indicated in the legend. The gray shaded area reflects able-bodied control’s Pre – Post (6m) data (95% percentile interval). Training the classifier on the pre-amputation data and testing it on the post-amputation data (and vice versus) revealed significantly above chance classification accuracies for all case-study participants at all post- amputation sessions (one-sample t-test: P1: Pre/1.5y: 89%; p<0.001; P2: Pre/5y: 67%; p<0.001; P3: Pre/6m: 88%; p<0.001). All other annotations are depicted in Figure 1.

Notably, this stability cannot be attributed to a pre-existing baseline difference, as hand activity pre-amputation was normal relative to controls (Supp. Figure 2A).

Similar pre-post stability was observed in motor cortex (M1; Supp. Figure 3A) and for the intact (unaffected) hand (Supp. Figure 4A).

Next, we investigated S1 finger representation stability in greater detail using a multi-voxel pattern analysis (Figure 2B; Methods). Multi-voxel activity patterns for the pre-amputated versus phantom fingers were significantly correlated at 6 months [five Pearson correlations per participant; P1: 0.68≤ *r*≤ .90, *p_uncorr_<*0.001; P2: 0.80≤*r*≤.85, *p_uncorr_<*0.001; P3: 0.88≤*r*≤.91, *p_uncorr_<*0.001]. Correlation coefficients at 6 months fell within the typical distribution seen in controls (see Supp. Figure 5 and Supp. Table 2 for control values). Similar stability was observed in M1 (Supp. Figure 3) and for the intact hand (Supp. Figure 5).

Combined, this confirmed that activity was largely stable before and after amputation at the single voxel level.

We next considered finger selectivity, i.e. activity profiles for each finger versus other fingers. Qualitative finger mapping revealed preserved somatotopy before and after amputation (Figure 2C). We applied a multivoxel pattern analysis using a linear support vector machine classifier (Figure 2D) to explore whether a pre-amputation-trained classifier can decode phantom finger movements (and vice versa). This analysis revealed significantly above chance classification for all case-study participants across all post-amputation sessions [Figure 2D; 2-3 one-sample t-tests per participant: P1 (Pre/1.5y): 90%; t(9)=10.5, *p_uncorr_<*0.001; P2 (Pre/5y): 67%; t(9)=4.85, *p_uncorr_<*0.001; P3 (Pre/6m): 89%; t(9)=11.0, *p_uncorr_<*0.001], with similar evidence in M1 (Supp. Figure 3).

We next investigated whether amputation reduces finger selective information, as suggested by previous cross-sectional studies^22^. Assessing for abnormalities in the pre-amputation data, we noted that 1 of the case-study participants, P2, exhibited lower classification for the pre-amputated hand relative to controls (Supp. Figure 2), likely due to P2’s impaired motor control pre-amputation (Supp. Video 1). Our key question remains whether this information degrades further following amputation. When comparing selectivity differences over 6 months relative to controls, none of the case-study participants showed significant reductions in average finger selectivity (Crawford t-test: P1: t(15)=-0.34, *p*=0.73; P2: t(15)=-0.24, *p*=0.80; P3: t(15)=-1.0, *p*=0.33; Supp. Figure 6C). While finger selectivity was reduced at P2 and P3’s final scan relative to their baseline (Figure 2D; 3 Wilcoxon tests per participant: P1 (1.5y): W=3.0, *p_uncorr_*=0.11; P2 (5y): W=2.0, *p_uncorr_*=0.005; P3 (6m): W=1.0, *p_uncorr_*=0.01), these reductions could be attributed to the much greater longitudinal variability between training and testing classifier samples^23^. Therefore, any reductions in finger selectivity could not be directly attributed to the amputation.

We also performed a complementary representational similarity analysis (RSA) using Mahalanobis distances (a continuous measure of finger selectivity), cross-validated across sessions. Similar to the decoding, RSA confirmed finger-selective information was significantly consistent across amputation for all case-study participants at all post-amputation timepoints (2-3 one-sample t-tests per participant: *p_uncorr_*<0.0001; Supp. Figure 6), with similar evidence in M1 (Supp. Figure 3C). We noted a few temporary, idiosyncratic (uncorrected) instances of reduced finger selectivity, relative to controls (Supp Figure 6). Using the RSA distances, we also tested the typicality of the inter-finger representational structure, an additional feature of hand representation. Correlating each participant’s inter-finger pattern to a canonical pattern revealed no deterioration in typicality scores 6-months post-amputation, compared to controls, with P3 even showing higher typicality than the average control (Crawford t-test: P1: t(15)=-0.9, *p*=0.38; P2: t(15)=-0.9, *p*=0.38; P3: t(15) = -3.5, *p*=0.003; Supp. Figure 6). Therefore, despite idiosyncratic reductions in finger selectivity, the representational structure was preserved post-amputation.

Finally, we examined changes in the lip representation, previously implicated with reorganization following arm amputation^4,7^. Projecting hand and lip univariate activity onto the S1 segments revealed no evidence of lip activity shifting into the hand region post-amputation (Figure 3A). All case-study participants showed typical longitudinal variability at their 6 months scan, relative to controls, for lip CoG [Figure 3B; Crawford t-test: P1: t(15)=0.25, *p*=0.80; P2: t(15)=-0.89, *p*=0.38; P3: t(15)=-0.9, *p*=0.37]. Further, lip activity in the S1 hand region at the final scan was typical [Figure 3C; P1 (1.5y): t(15)=0.8, *p*=0.20; P2 (5y): t(15)=-0.5, *p*=0.71; P3 (6m): t(15)=1.2, *p*=0.10]. Also, when visualizing the lip map boundaries within S1 for all sessions, using a common minimum threshold, there was no evidence for an extension of the lip map (Figure 3D). Examining multivariate lip representational content, P2 showed an increased lip-to-thumb multivariate distance at their 6 months scan, relative to controls [Figure 3E; Crawford t-test: P1: t(15)=0.69, *p*=0.25; P2: t(15)=3.1, *p*=0.003; P3: t(15)=.74, *p*=0.23; intact hand and feet data included in Supp. Figure 7] However, it returned to the typical range of controls when assessed at their 5-year timepoint. Similar stability was found in M1 (Supp. Figure 3), and the unaffected hemisphere (Supp. Figure 4). These results demonstrate that amputation does not affect lip topography or representational content in S1.

**Figure 3.**
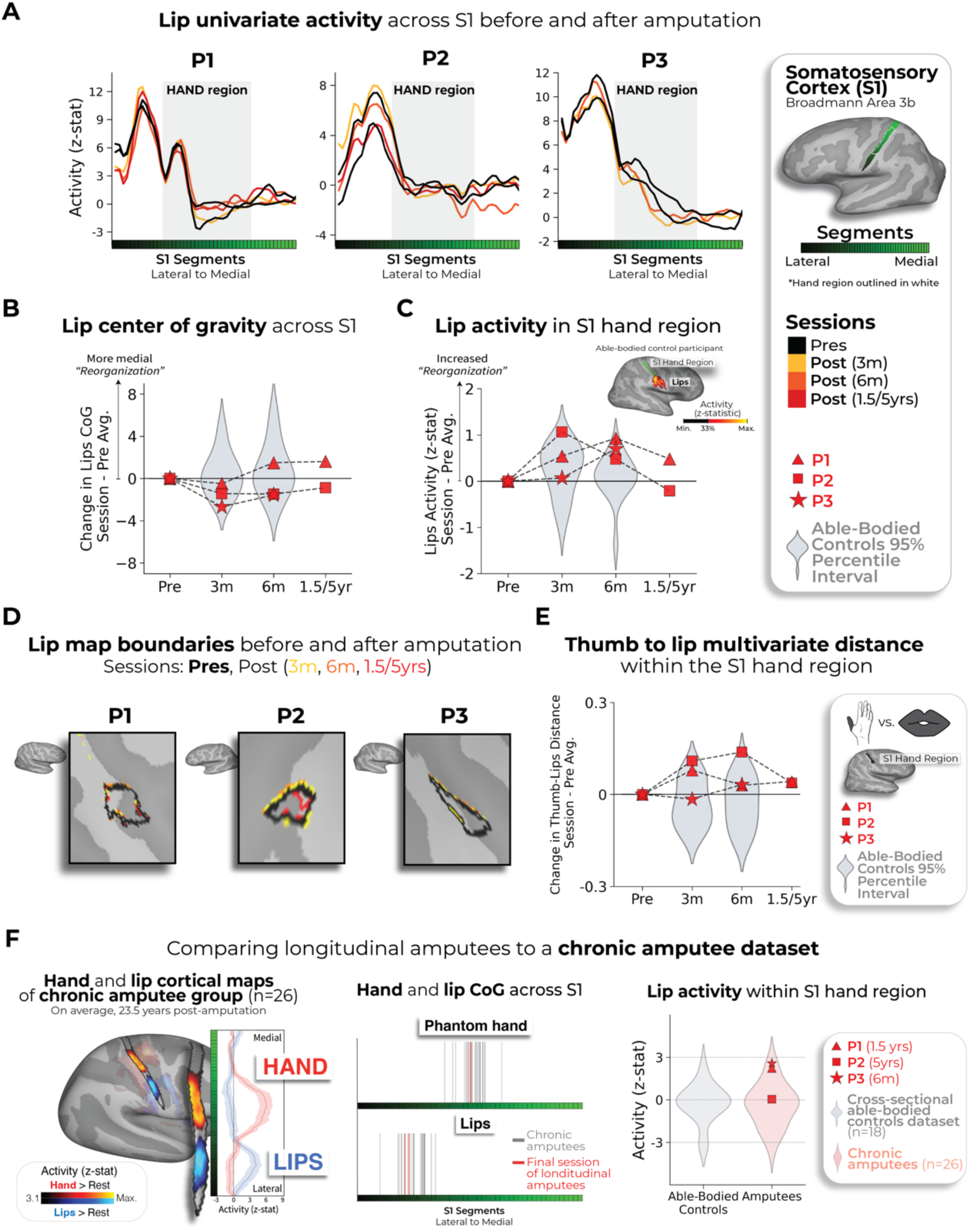
No evidence for lip reorganization after amputation. **(A)** Each case-study participant’s lip activity (versus rest) for their sessions projected across the S1 ROI. Black lines reflect pre-amputation activity, yellow (3m), orange (6m) and red (1.5/5y) lines post-amputation. Grey region depicts approximated coverage of the hand portion within S1. **(B)** All case-study participants showed typical longitudinal variability at their 6 months scan, relative to controls, for lip CoG. Positive values reflect medial shifts (towards the hand). **(C)** All case-study participants showed typical lip activity in the S1 hand region at the final scan. Right corner of panel depicts representative control participant’s activity for the hand and lips (versus feet; minimally thresholded at 33% the max. z-statistic). **(D)** All case-study participants exhibited no expansions of the lip map boundaries towards the hand region. Maps masked to the S1 ROI and minimally thresholded (Z > 4.5). **(E)** All case-study participants showed stable thumb-to-lip multivariate Mahalanobis distances cross-validated at their final scan, relative to controls. **(F)** Comparing the case-study participants to a chronic amputee dataset (n=26). **Left** – chronic amputee’s group-level cortical activation maps of the phantom hand and lips (versus rest) projected onto a single hemisphere (minimally thresholded at Z > 3.1). Opacity applied to activity outside the S1 ROI. Group univariate activity plotted as a line (group mean ± standard error) for the phantom hand (red) and lips (blue) across the S1 ROI. **Middle** – All case-study participants, comparable to chronic amputees, showed a typical center of gravity for both the phantom hand (top row) and lips (bottom row). **Right** – All case-study participants exhibited typical lip activity within the S1 hand region during their final session consistent with chronic amputees. The magnitude of lip activity (95% percentile interval) within the S1 hand region for a secondary able-bodied control group (n=18; shown in grey). Chronic amputees shown in pink and the case-study participants last session data shown in red. All other annotations are the same as described in Figure 2.

To complement our longitudinal findings, we compared our case studies to a cohort of 26 chronic upper-limb amputee participants, on average 23.5 years post-amputation (Figure 3F; individual hand and lip cortical maps in Supp. Figure 8). Our case-studies’ topographical features were comparable to chronic amputees for both the phantom hand [Crawford t-test: P1 (1.5y): t(15)=0.28, *p*=0.77; P2 (5y): t(15)=0.29, *p*=0.77;*p*=0.77; P3 (6m): t(15)=0.28, *p*=0.22; *p*=0.82] and lips [P1 (1.5y): t(15)=0.53, *p*=0.59; P2 (5y): t(15)=0.01, *p*=0.98; P3 (6m): t(15)=0.37, *p*=0.71]. Average lip activity within the S1 hand region was slightly (though not significantly) higher for a few of our case-studies relative to chronic amputees (Crawford t-test: P1 (1.5y): t(15)=1.6, *p*=0.10; P2 (5y): t(15)=0.24, *p*=0.81; P3 (6m): t(15)=1.8, *p*=0.065), reflecting that lip activity does not steadily increase in the years after amputation. Collectively, these results provide long- term evidence for the stability of hand and lip representations despite amputation.

Beyond the stability of the lip representation across amputation, our findings reveal highly consistent hand activity despite amputation. This unchanged hand representation challenges the foundational assumption that S1 activity is primarily tied to peripheral inputs, suggesting that S1 is not a passive relay of peripheral input, but an active supporter of a resilient ‘model’ of the body—even after amputation. We therefore conclude that, in the adult brain, S1 representation can be maintained by top-down (e.g. efferent) inputs. This interpretation sheds new light on previous studies showing similar S1 topographical patterns activated by touch^24^, executed movement^25^ and planned movement^26^.

Due to the limitations of non-human models that cannot communicate phantom sensations, it is not surprising that the persistent representation of a body part, despite amputation, has been neglected from previous studies. Without access to this subjective dimension, researchers may have missed the profound resilience of cortical representations. Instead, previous studies determined S1 topography by applying a ‘winner takes all’ strategy –– probing responses to remaining body parts and noting the most responsive body part in the input-deprived cortex^3,4^.

Ignoring phantom representations in these analyses leads to severe biases in the interpretation of the area’s inputs (as demonstrated in Supp. Figure 10).

Combined with cross-sectional designs, this has incorrectly led to the impression of large-scale reorganization of the lip representation following amputation. Our longitudinal approach reveals no signs of reorganization in S1—not even subtle upregulation from homeostasis—further reinforcing the notion that S1 is not governed by deprivation-driven plasticity.

For brain-computer interfaces, our findings demonstrate a highly detailed and stable representation of the amputated limb for long-term applications^27^. For phantom limb pain treatments, our study indicates that targeted muscle reinnervation and regenerative peripheral nerve interfaces do not ‘reverse’ reorganization or alter the cortical hand representation^22,28^. Finally, our findings affirm the unaltered nature of adult sensory body maps following amputation, suggesting Hebbian and homeostatic deprivation-driven plasticity is even more marginal than considered by even the field’s strongest opponents of large-scale reorganization^17,29^.

## Supplementary Results

**Supplementary Figure 1.**
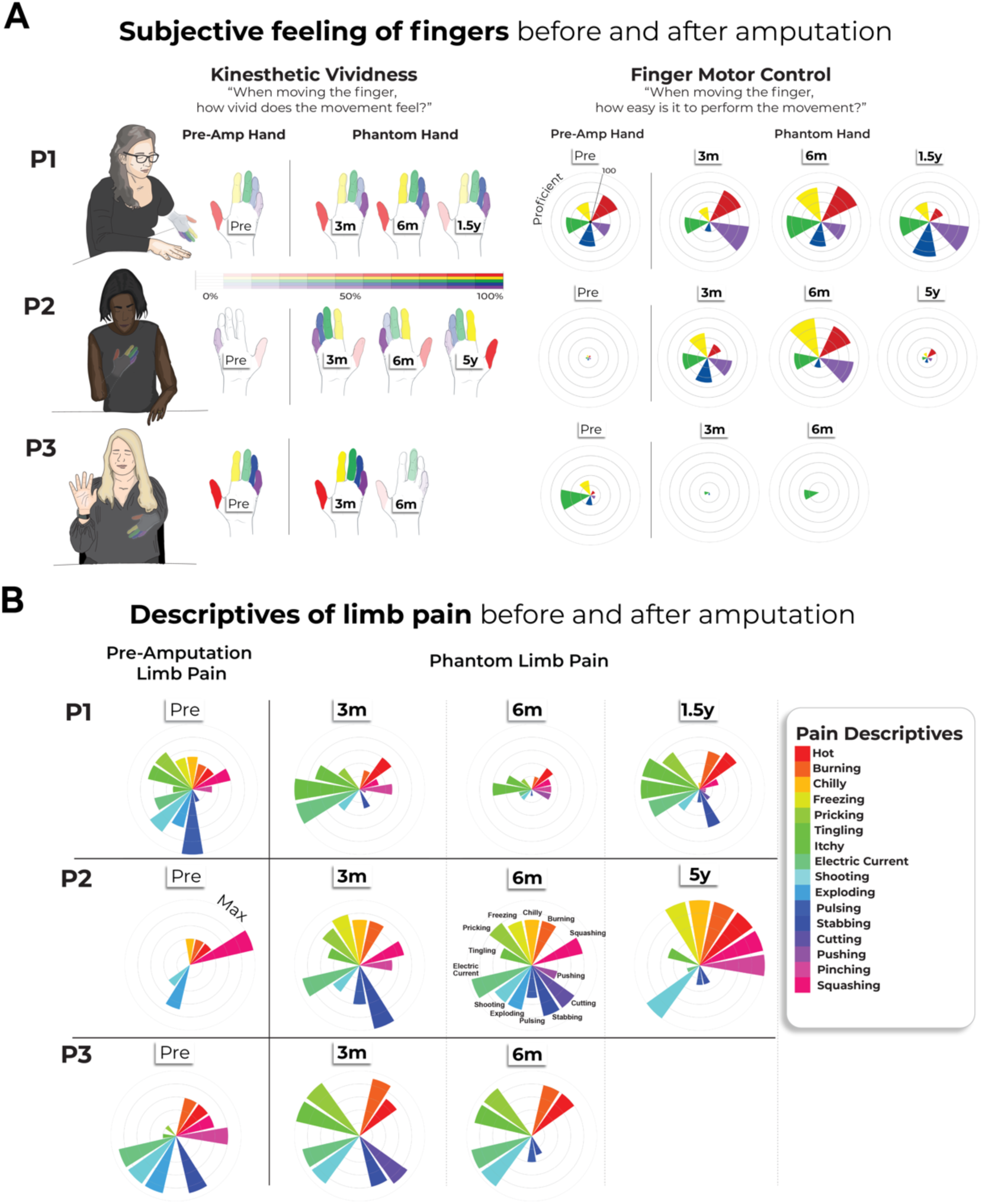
Longitudinal characterization of finger sensations and limb pain. **(A)** Affected hand sensations before and after amputation. Finger vividness and motor control for the phantom fingers, relative to the pre-amputated fingers. Kinesthetic vividness rated on a scale from 0 (no sensation) to 100 (as vivid as the unaffected hand) with color intensity indicating level. Movement difficulty rated from 100 (as easy as the unimpaired hand) to 0 (extremely difficult). Finger colors: red=D1, yellow=D2, green=D3, blue=D4, purple=D5 (palm excluded). **(B)** Before and after amputation, participants reported intensity values for each pain descriptive word, broadly categorized into sensations that are mechanical, temperature-related and other. For each word, participants were asked to describe the intensity between 0 (non-existing) to 100 (excruciating pain) as it relates to that particular word. A value of 100 (Max) is the largest radii on the polar plot. 3M=3months post-amputation; 6M=6months post- amputation. 1.5/5yrs=1.5 or 5 years post-amputation.

**Supplementary Figure 2.**
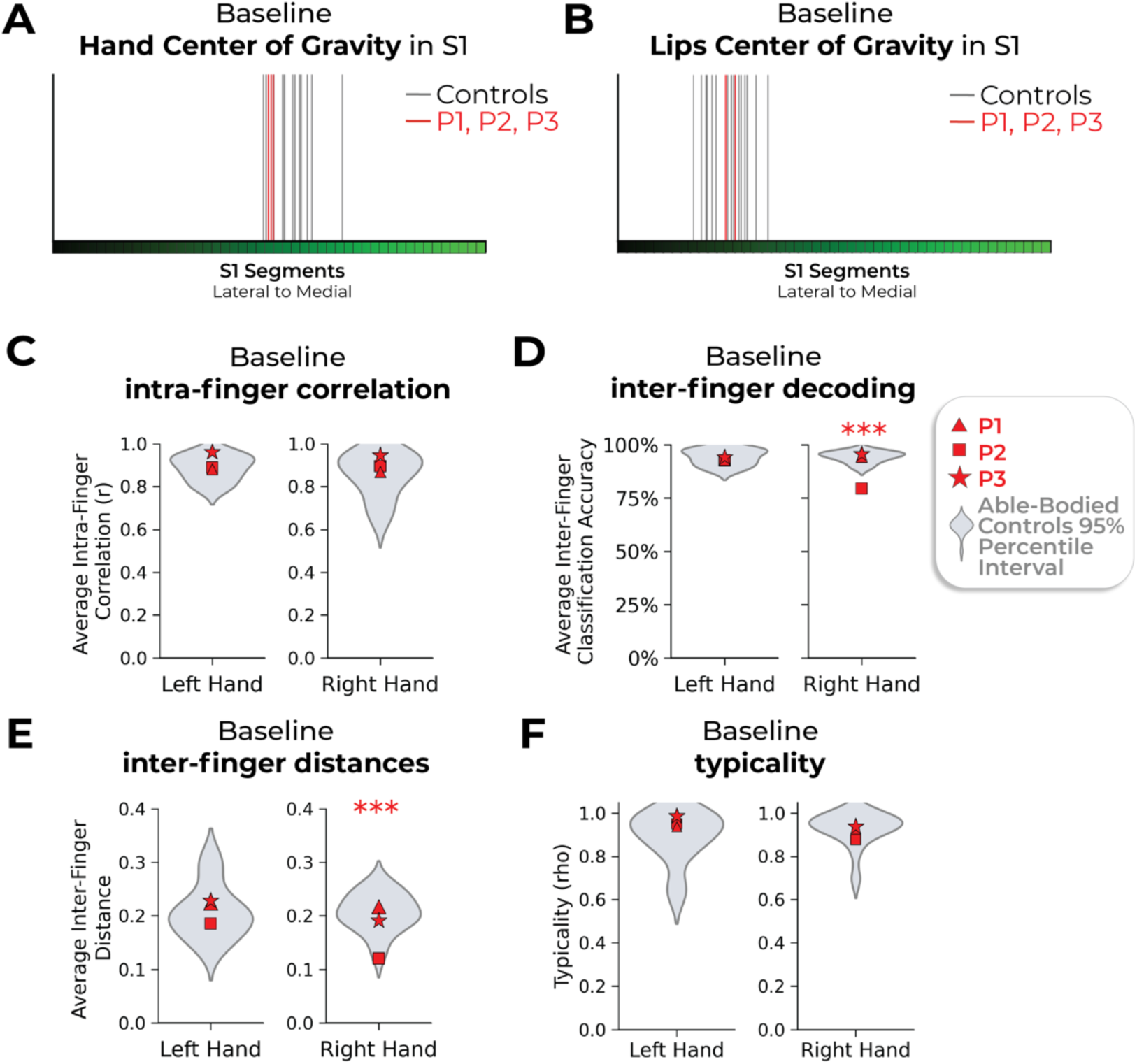
Baseline measures for the case-study participants that underwent an amputation versus able-bodied controls. Across all panels, we only report statistics when significant. Case-study participants showed similar responses to able-bodied controls in the baseline (pre-amputation) S1 center of gravity for the **(A)** hand and **(B)** lips. **(C)** All case-study participants had similar average intra-finger correlations between the two pre-sessions as controls. For baseline average inter-finger **(D)** classification accuracy and **(E)** distances. One case-study participant exhibited lower values for their affected hand only, relative to controls [Crawford t-test: decoding and distances: P2: p<0.001] **(F)** All case-study participants had similar hand typicality between the two pre-sessions as controls. All other annotations the same as described in Figures 2 *and 3*.

**Supplementary Figure 3.**
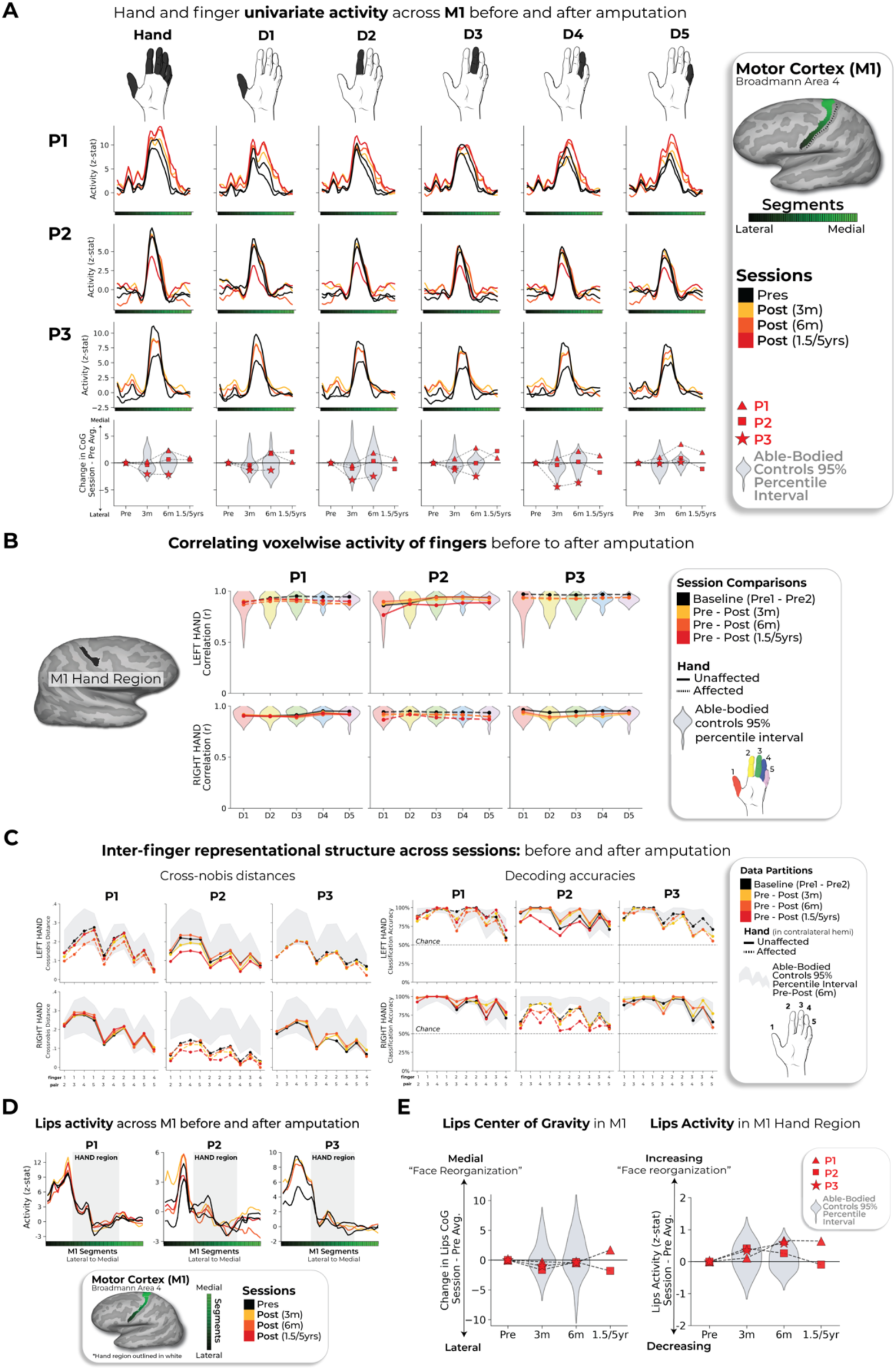
Replication of all primary results within motor cortex. Across all panels, we only report statistics when significant. **(A)** Hand and finger univariate activity across M1 before and after amputation. When testing the stability of the whole hand condition across sessions, all case-studies fell within the distribution of controls at all timepoints. **(B)** When correlating voxel wise finger activity across sessions, all case-studies exhibiting similar correlation coefficients as controls, for all fingers. Please refer to the Supp. Figure 5 caption for a more detailed understanding of the correlation analysis. **(C)** Inter-finger representational structure across sessions, measured using cross-nobis distances (left) and decoding accuracies (right). First, when assessing for atypicality in our case-studies pre-amputation compared to controls, only case-study P2 exhibited reduced average finger selectivity pre-amputation based on the RSA (Crawford t-test: t(15)=-3.15, p=0.007) and decoding (t(15)=-3.9, p=0.001; similar to what was observed in S1). Next, when testing for reductions in average finger selectivity at the 6-month timepoint, relative to baseline, only case-study P1 exhibited a significant reduction compared to controls [cross-nobis distances: 3 comparisons; t(15)=2.33; p_uncorr_=0.02); decoding: 3 comparisons; t(15)=2.32; p_uncorr_=0.03]. However, it returned to the typical range when later assessed at the 1.5 year timepoint (for both measures). We also noted that case-study P3 showed a significant reduction at the 6-month timepoint, relative to controls, in the decoding (3 comparisons; t(15)=2.18, p_uncorr_=0.046), but not the cross-nobis. **(D)** Lips univariate activity plotted across M1 before and after amputation. **(E)** All case studies showed typical session to session variability as controls in (left side) the lips center of gravity across M1 and (right side) lips activity in the M1 hand region. All annotations are the same as described in the captions of the Figures 2-3 and Supp. Figure 5.

**Supplementary Figure 4.**
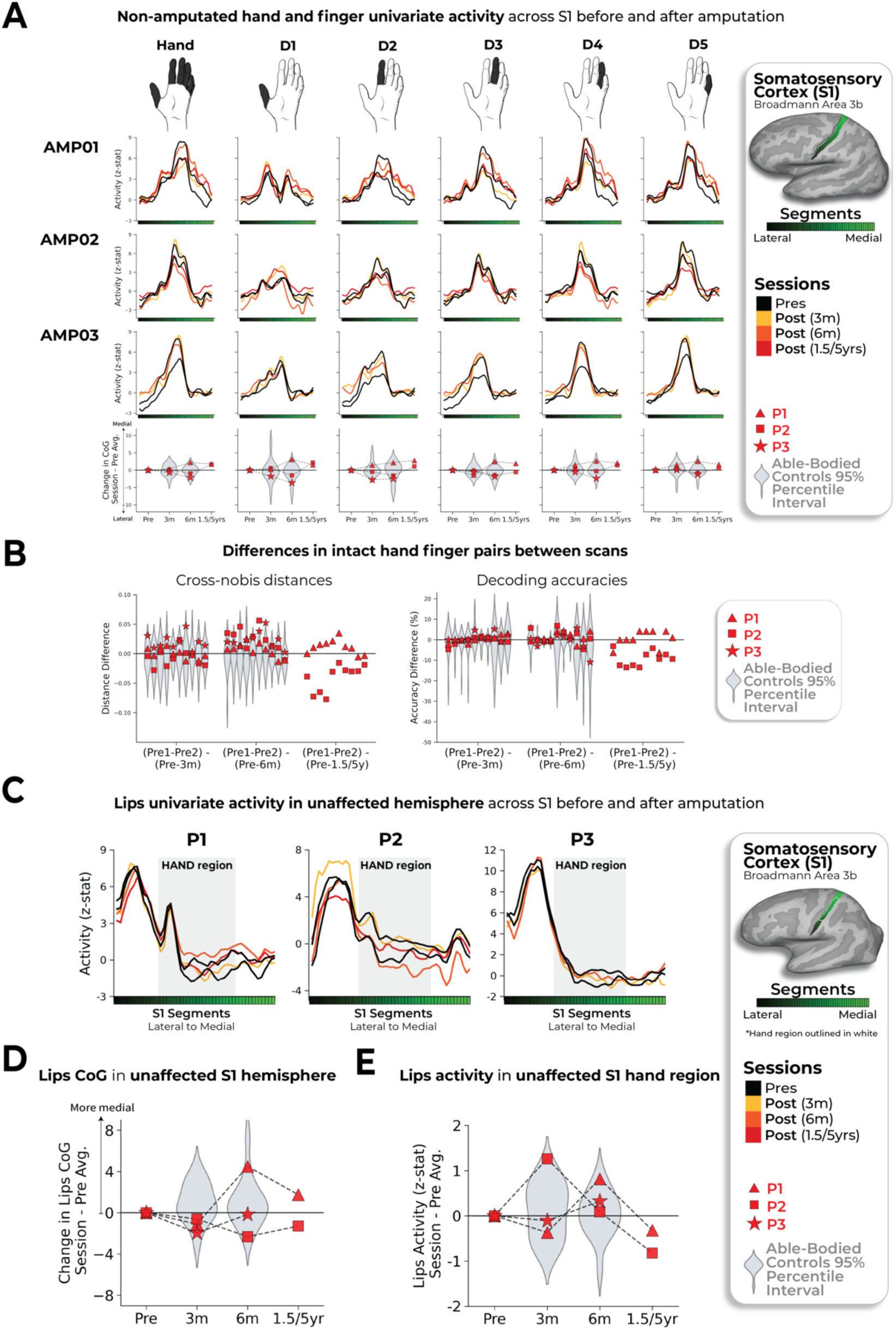
Stability of the intact (non-amputated) hand and lip topography in the non-affected hemisphere across amputation. **(A)** Intact hand and finger univariate activity across S1 before and after amputation. When testing the stability of the whole hand condition across sessions, all case-studies fell within the distribution of controls at all timepoints. **(B)** Unaffected (intact) hand between-session differences in inter-finger values. Difference values are depicted for the (left) cross-validated distances and (right) decoding accuracies. Classification/distance differences before and after amputation are visualized for each finger pair [Pre1-Pre2] minus [Pre Avg. – Post1 (3m)] minus, [Pre1-Pre2] minus [Pre Avg. – Post2 (6m)] and [Pre1-Pre2] minus [Pre Avg. – Post3 (1.55/y)]. Each violin plot reflects an individual finger pair (same order of finger-pairs as detailed in Figure 2D). For consistency, the control values are all for the left-hand. When computing the session-to-session differences relative to controls, all case-study participants showed typical session-to-session variability in finger selectivity at the 6-month timepoint, relative to controls. **(C)** Longitudinal lips univariate in the unaffected hemisphere (contralateral to intact hand) across S1 before and after amputation. **(D)** All case study participants showed typical changes in the lips center of gravity (CoG) in the unaffected S1 hemisphere across scans, relative to controls. **(E)** When testing for changes in lip activity (in the unaffected hand region), one case-study, P1, exhibited a significant atypical increase in lip activity relative to controls at the 6-month timepoint (Crawford t-test: t(15)=2.75, p_uncorr_=0.01). However, the activity returned into the distribution of controls when tested at the 1.5 year timepoint (t(15)=0, p_uncorr_=0.99). All other annotations are the same as described in Figures 2 *and 3*. We only report statistics when significant.

**Supplementary Figure 5.**
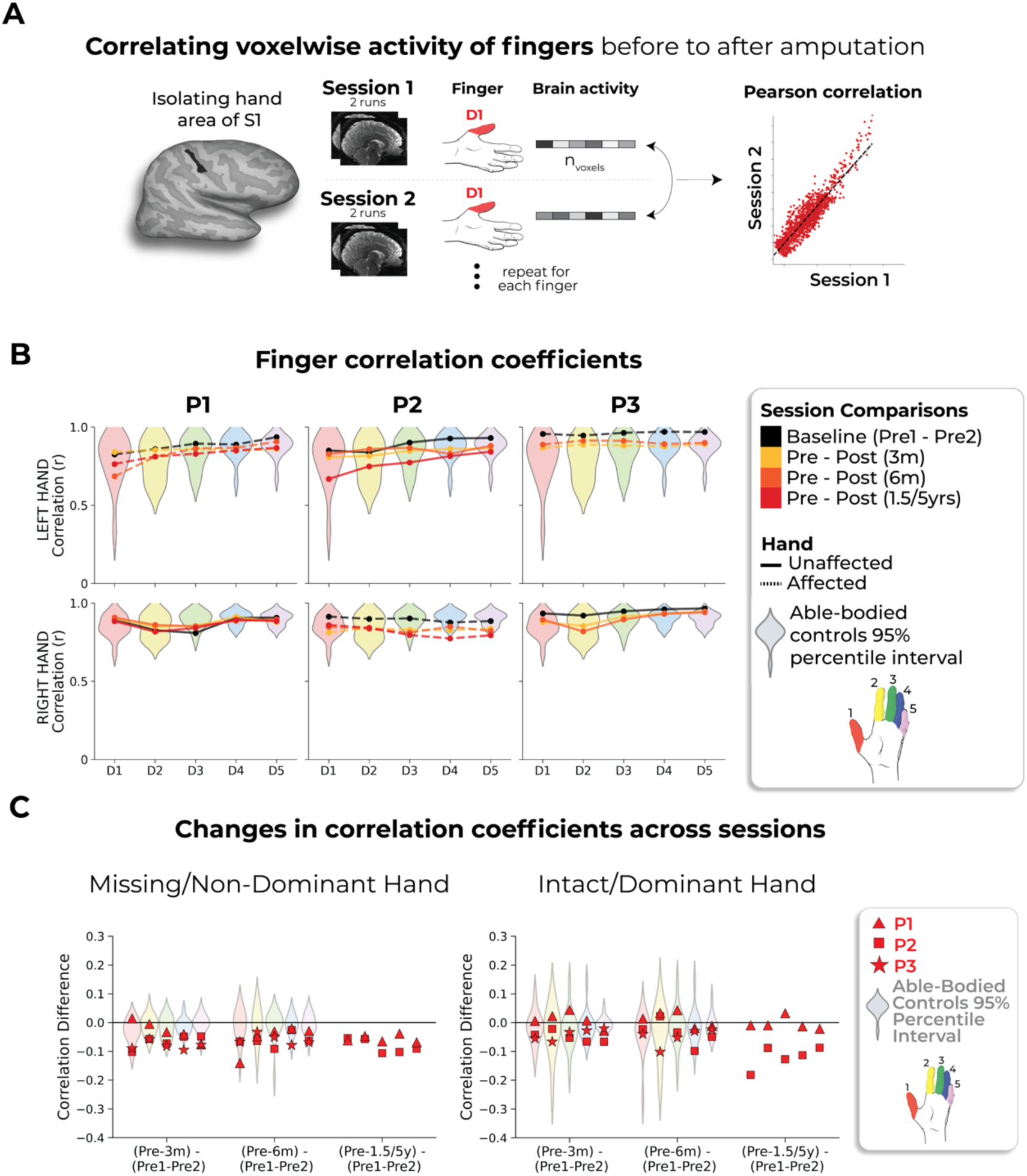
Correlating pre- to post-amputation multivoxel finger activity patterns. **(A)** Visualization depicting the inter-session Pearson correlations of individual fingers within the BA3b hand region. **(B)** Inter-session correlations for the left (top row) and right hands (bottom) in the contralateral hand ROI. Line colors indicate session pairings (indicated in the legend). For case-study participants, dashed line denotes the affected hand; solid line unaffected hand. Violin plots reflect able-bodied control’s Pre – Post (6m) values. **(C)** Between-session differences in finger correlation coefficients. Difference values are depicted for the (left) missing or non-dominant hand of controls and (right) intact or dominant hand of controls. The difference values are ordered to reflect the increasing gap between sessions: [Pre1-Pre2] minus [Pre Avg. – Post1 (3m)] minus, [Pre1-Pre2] minus [Pre Avg. – Post2 (6m)] and [Pre1-Pre2] minus [Pre Avg. – Post3 (1.55/y)]. Each violin plot reflects an individual finger. When testing whether the case-study participants showed a unique reduction in the average correlation, across fingers, relative to controls, for the missing hand, only P3, at the 3-month timepoint, for the missing hand (not intact), showed a significant pre-post reduction in the average correlation coefficient, relative to controls (t(15)=-2.59, p_uncorr_=0.02). However, this difference returned to the typical range of controls when later tested at the 6-month timepoint (t(15)=-1.23, p_uncorr_=0.23). All other annotations are as in Figure 2. We only report statistics when significant.

**Supplementary Figure 6.**
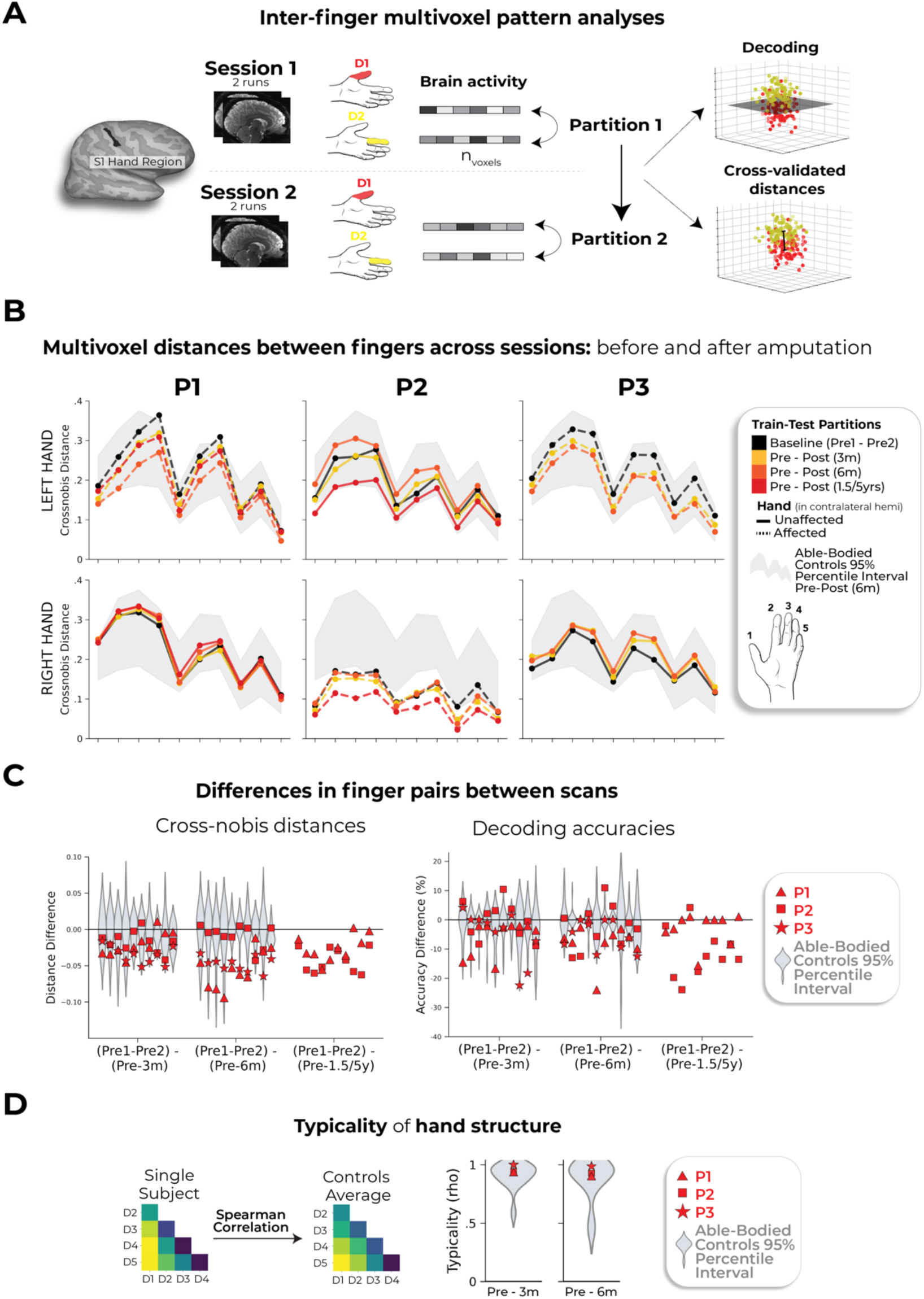
Representational similarity analysis of inter-finger representational structure. **(A)** Graphic illustration of multivoxel pattern analyses. **(B)** Inter-finger multivariate analysis using cross-validated Mahalanobis (cross-nobis) distances. Line colors denote train-test/cross validation session pairs, respectively as indicated in the legend. The gray shaded area reflects able bodied control’s Pre – Post (6m) data (95% percentile interval). **(C)** Classification/distance differences before and after amputation are visualized for each finger pair [Pre1-Pre2] minus [Pre Avg. – Post1 (3m)] minus, [Pre1-Pre2] minus [Pre Avg. – Post2 (6m)] and [Pre1-Pre2] minus [Pre Avg. – Post3 (1.55/y)]. Each violin plot reflects an individual finger pair (same order of finger-pairs as detailed in B). When comparing differences relative to controls, we observed some temporary, idiosyncratic reductions in average finger selectivity, relative to controls. First for the cross-nobis results, P1 showed a temporary reduction in average finger selectivity at 6 months (3 comparisons; t(15)=-2.79, p_uncorr_=0.01), though later offset to the typical range at their follow-up 1.5-year scan. P2 only exhibited reduced selectivity only at the 5-year timepoint, though reduction seen in the intact hand as well (Supp Figure 4). Finally, P3 exhibited reduced selectivity at 6 months relative to controls (2 comparisons; t(15)=-2.36, p_uncorr_=0.03). For the decoding results, P2 seemed to show significantly reduced selectivity at the 5-year timepoint, though also reduced for the intact hand (Supp Figure 4). **(D)** The representational typicality of the hand structure was estimated by correlating each session’s cross-validated Mahalanobis distances for each participant to a canonical inter-finger structure (controls average). All case-study participant’s typicality values fell within the distribution of controls. All other annotations are as in Figure 2. We only report statistics when significant.

**Supplementary Figure 7.**
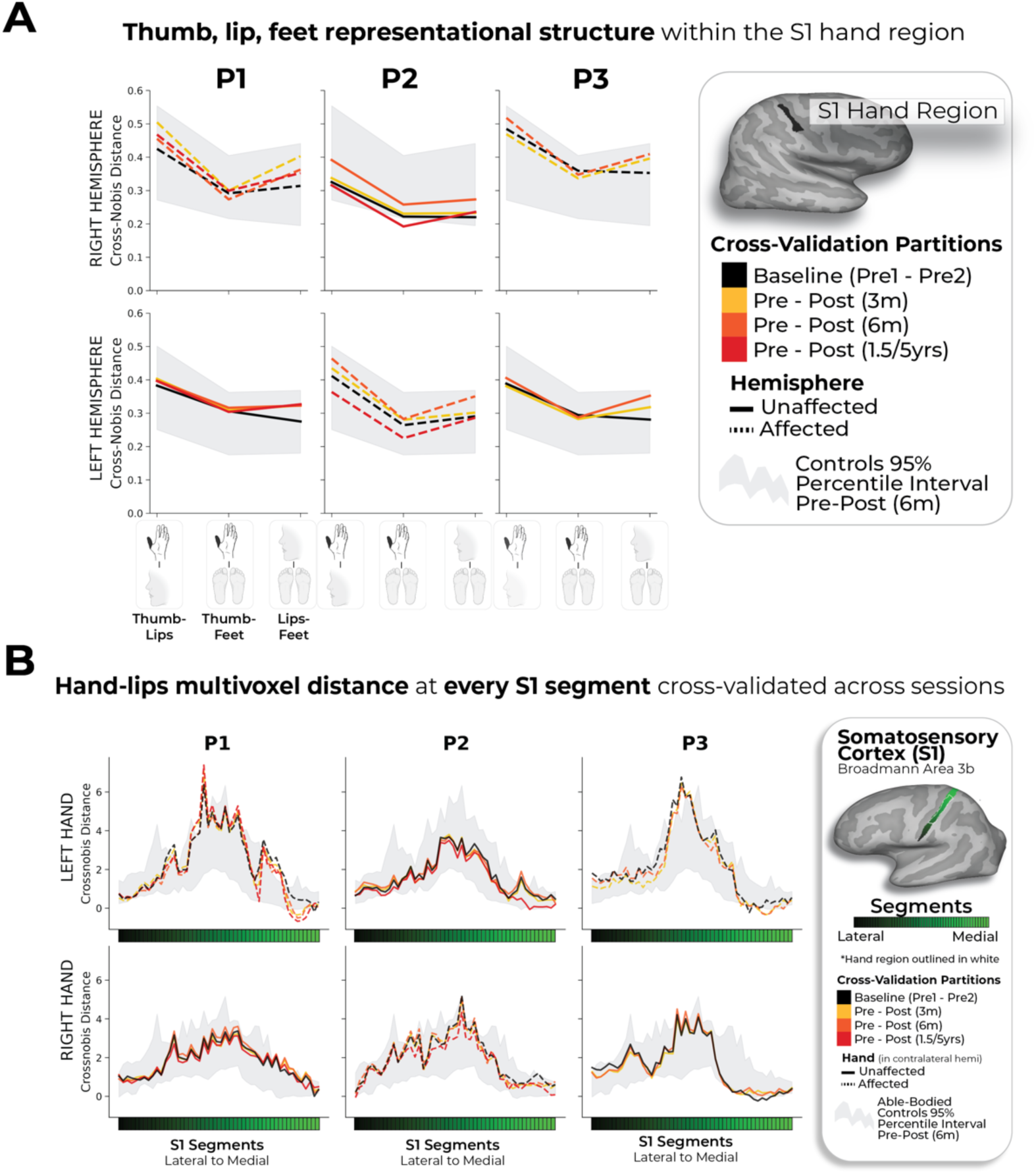
Thumb, lip and feet distances within the S1 hand region. **(A)** Multivariate distances between the thumb, lip and feet cross-validated across sessions depicted for the right (top row) and left hemisphere (bottom) of the case-study participants that underwent an amputation and controls, contralateral to the thumb side being moved. Distances appear in the following order: (1) thumb-lips, (2) thumb-feet, (3) lips-feet. Line colors indicate session pairings (indicated in the legend). For case-study participants, dashed line denotes the affected hemisphere; solid line unaffected hemisphere. Grey shaded area reflect able-bodied control’s Pre – Post (6m) values. For the affected hemisphere of the case-study participants, all distances fell within the typical range of the able-bodied controls. (B) We also tested whether changes occurred in the multivariate hand-lip distance when performed within each of the 49 S1 segments/ All case-study participants showed similar distances across sessions, before and after amputation. All other annotations are the same as described in Figure 2.

**Supplementary Figure 8.**
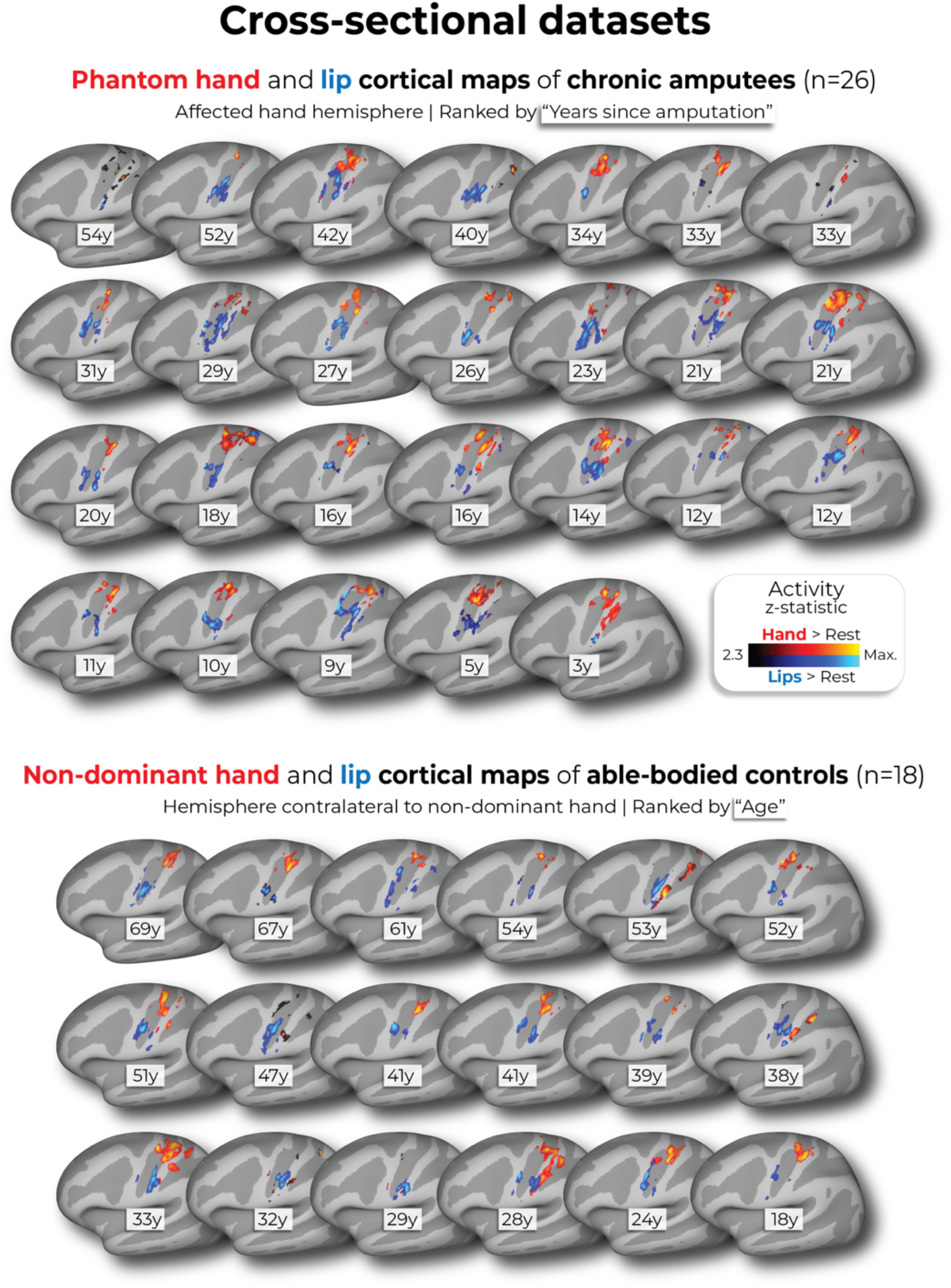
Hand and lip cortical maps of cross-sectional datasets. Participant hand and lip cortical maps – registered to a standard cortical surface – are visualized for the chronic amputee participants (top row; n=26) and secondary able-bodied control participants who underwent the same procedures as the chronic amputees (n=18; bottom row). Hand maps for the amputees reflect moving their phantom hand, while for controls reflect moving their non-dominant hand (in the contralateral hemisphere). All maps are contrasted against rest, minimally thresholded at 50% the maximum z-statistic and masked to Broadmann regions: 1, 2, 3a, 3b, and 4. Amputee maps are ranked by the numbers of years since amputation at the time of the scan and control maps are ranked by the participants age at the time of the scan. All other annotations are the same as described in Figure 1.

**Supplementary Figure 9.**
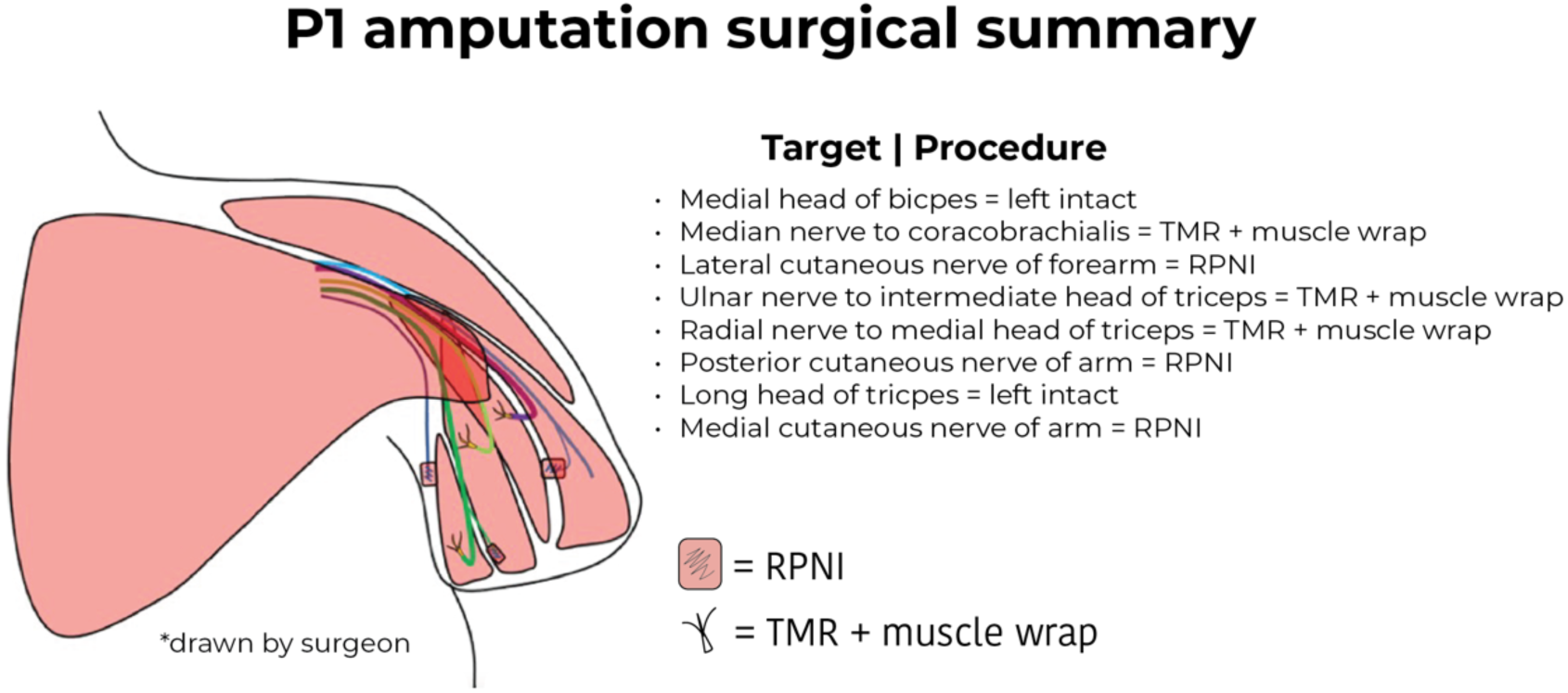
Summary of P1’s amputation procedure. An illustration depicting the unique amputation surgery of P1’s left arm, as well as summary of the procedures performed for each respective nerve. TMR=targeted muscle reinnervation; RPNI=regenerative peripheral nerve.

**Supplementary Figure 10.**
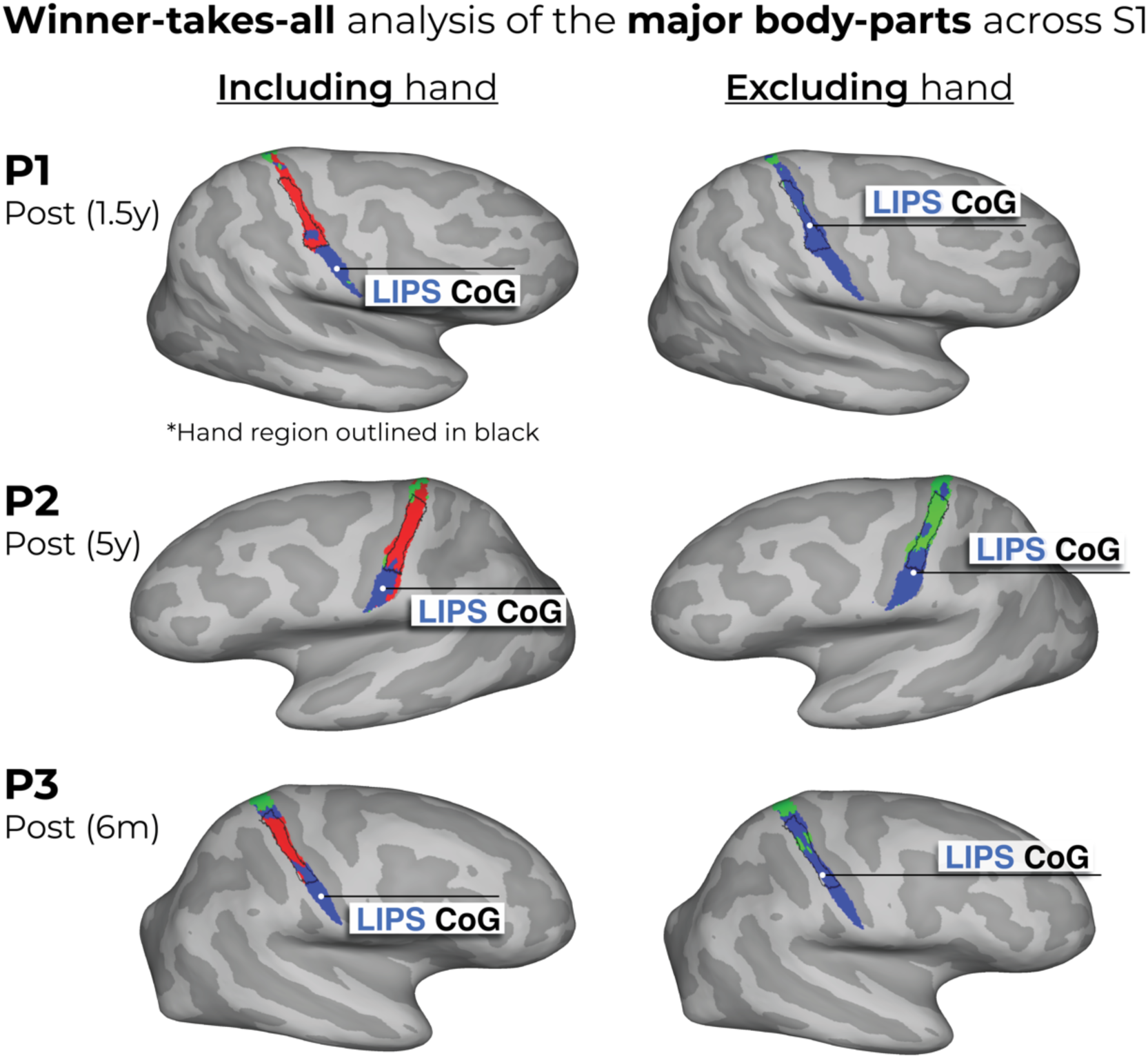
Winner-takes-all analysis of the major body parts (hand, lips and feet) across S1. Using the data from the last session of each participant, each voxel was awarded to the body-part with the highest response. Left column – we show the winner-takes-all analysis when performed on 3 body-parts: hand (red), lips (blue) and feet (green) versus (Right column) when excluding the physically absent hand. This comparison reveals supposed large-scale expansions of the lips or feet into the deprived hand region (black outline) post-amputation. We’ve also depicted the center of gravity (CoG) of the winner-takes-all lip cluster (white circles) to further demonstrate this. When excluding the hand activity, the CoG of the lips ‘shifts’ towards the hand area. Thus, ignoring the primary body part – depending on your analysis choices – can substantially bias the results^30,31^. Combined with the use of cross-sectional designs, this analysis approach has led to the impression of cortical remapping and even large-scale reorganization of the lip representation following amputation. Crucially, the newly assigned winner in the hand area [left panel] has rarely been directly compared against the persistent representation of the missing hand, and indeed, indicative evidence show that this recorded activity in the hand area is weak (we extensively discuss this in our recent review ref.^17^).

**Supplementary Figure 11.**
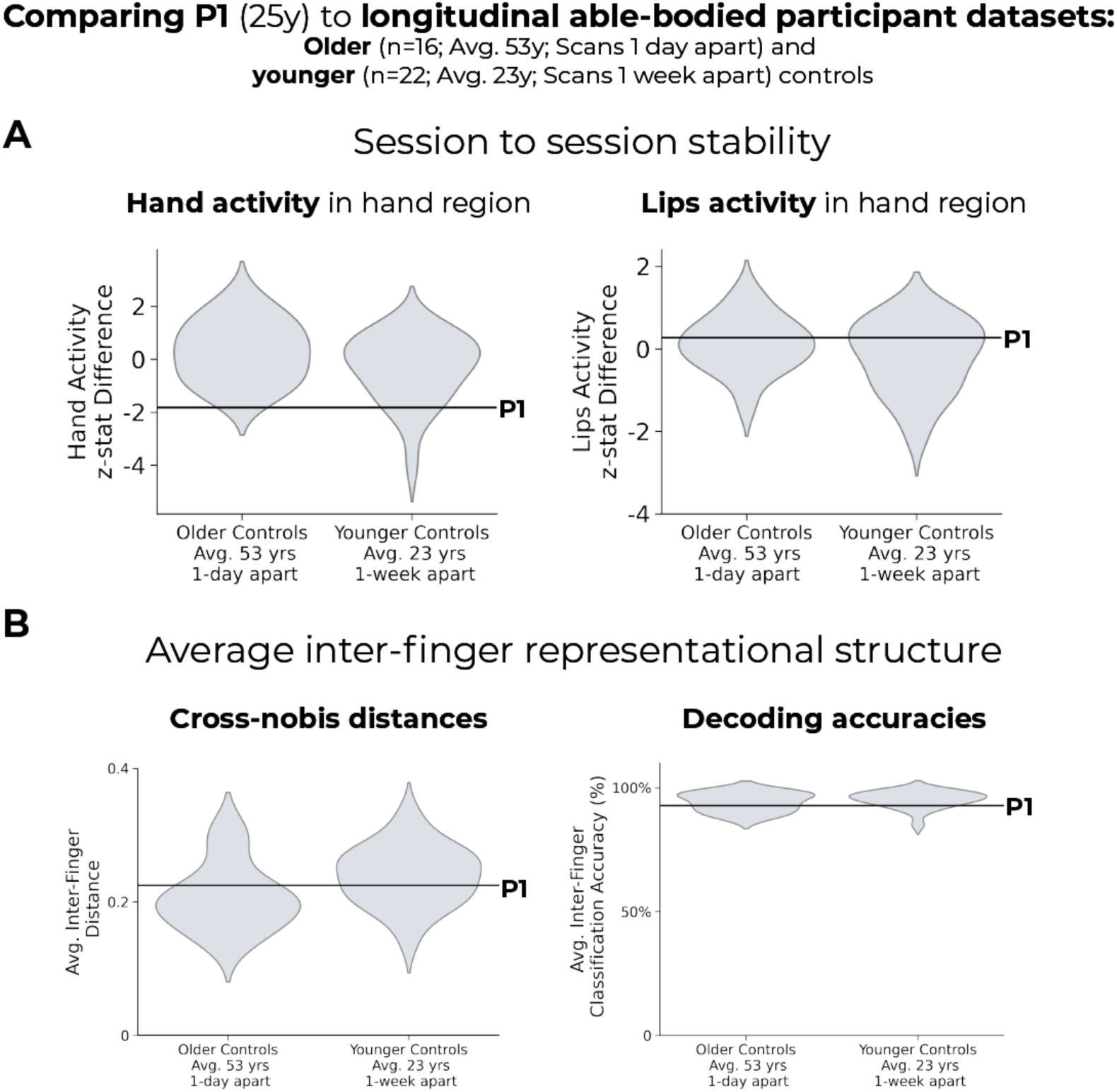
Comparing case study P1’s hand and lip activity to 2 longitudinal able-bodied participant datasets: older and younger controls. As the longitudinal able-bodied controls were age-matched to P2 and P3, we investigated whether younger able-bodied controls (see Methods) showed significant differences on multiple measures compared to the older controls. The older controls are the longitudinal controls described in the main text. The younger controls performed the same task on the same scanner, 2 scans separated by 1-week. We examined the session-to-session stability in our primary univariate and multivariate measures. First, for the univariate measures, younger and older controls showed similar session to session differences in **(A)** hand activity in the S1 hand region (independent samples t-test: t(37)=1.7, p=0.09), and **(B)** lip activity in the hand region (t(37)=1.3, p=0.18). Further, P1’s pre-amputation scan data (black line) showed no significant difference between older controls and younger controls for either body-part (hand: P1 vs. Older: t(15)=-1.7, p=0.09; P1 vs. Younger: t(21)=-1.1, p=0.2; lips: P1 vs. Older: t(15)=0.2, p=0.8; P1 vs. Younger: t(21)=0.52, p=0.6). Grey violin plots reflect controls data (95% percentile interval). **(C)** Next, for the multivariate measures, younger controls showed a trend for higher average inter-finger representational structure compared to older controls, in the cross-nobis distances (t(37)=-1.95, p=0.06), but not the decoding (t(37)=-0.87, p=0.38). P1’s pre-amputation session data was not different than the older or younger control groups for either measure (cross-nobis: P1 vs. Older: t(15)=0.32, p=0.75; P1 vs. Younger: t(21)=- 0.32, p=0.74; decoding: P1 vs. Older: t(15)=-0.35, p=0.72; P1 vs. Younger: t(21)=-0.70, p=0.48). All other annotations are the same as those described in Figure 2.

**Supplementary Figure 12.**
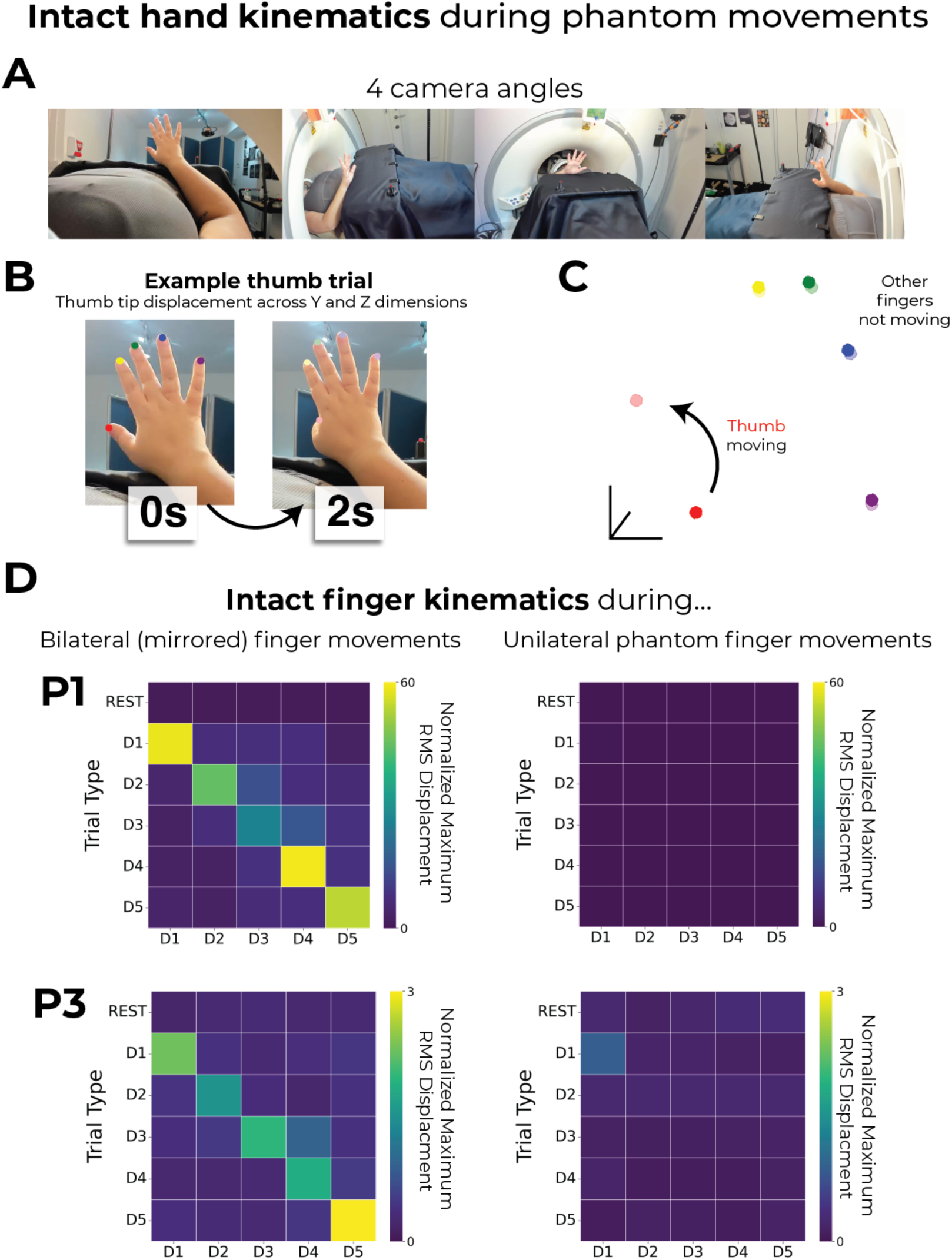
Intact finger kinematics during mirrored and phantom finger movements. **(A)** To test whether the intact fingers are being moved simultaneously during phantom finger movements, we tested 2 of the 3 case-study participants on a finger tapping task. Each participant was positioned inside an MRI scanner. We visually cued each participant to perform a finger flexion movement (each 2-seconds; 5 fingers or REST; 7 repetitions per condition). There were two blocks: bilateral (mirrored) finger movements, where participants were told to mirror the movements of the intact and phantom fingers, and unilateral phantom finger movements, where participants were told to move the phantom fingers. Participants were randomly cued which finger to move (or REST). We recorded kinematics of the intact fingers, using 4 cameras (Logitech brio, 1080p, 60fps). **(B-C)** Using Anipose’s triangulation function^32^ to triangulate the 4 cameras into 3D coordinates, we defined the 3D coordinates of the tip of each finger. Using the 3D coordinates, we then computed the root mean square (RMS) displacement of each dimension (x, y, z) within a trial. Across dimensions, we selected the dimension with the highest RMS displacement. We then averaged across repetitions of the same trial type. Finally, we normalize these values relative to the RMS displacement observed in the REST condition, effectively capturing relative movement magnitude. We provide a single trial visualization of each finger’s 3D coordinates (for the y and z dimensions) at the first (dark colours) and last (light colours) timepoints of a single move thumb trial. Note the distinct individuation of the thumb and not the other fingers. **(D)** We observed that while bilateral mirror finger movements show clear finger individuation of the intact fingers (plots on the left), the intact fingers do not move during phantom finger movements (plots on the right).

**Supplementary Table 1.** Demographics of case-study participants that underwent an amputation. PLS = phantom limb sensation; Limb pain reflects pre-amputation limb pain or post-amputation phantom limb pain. Frequency scores: 1 – all the time, 2 – daily, 3 – weekly, 4 – several times per month, and 5 – once or less per month. Chronic pain/sensation values were calculated by dividing intensity by frequency. NA = not available/applicable. Upper extremity functional index measures participant difficulty with performing activities due to their missing limb.

|  | **P1** | **P2** | **P3** |
| --- | --- | --- | --- |
| **Sex** | Female | Female | Female |
| **Age** (at first scan) | 26 | 57 | 49 |
| **Handedness at birth** | Left-handed | Right-handed | Right-handed |
| **Cause of amputation** | Arteriovenous vascular malformation (AVM) | Sarcoma tumour | Severell-Martorell syndrome led to multi-fractured arm with bones not healing |
| **Disability duration** | AVM progressed over a few years | Tumour slowly developing since 1995 | Musculoskeletal issues since childhood |
| **Amputated limb** | Left upper limb | Right upper limb | Left upper limb |
| **Level of amputation** | Transhumeral | At elbow | Transhumeral |
| **Amputation surgery** | Combination of targeted muscle reinnervation and regenerative peripheral nerve interfaces, see Supp. Fig. 9. | Traditional: sharply transected the nerves and allowed to retract | Traditional: sharply transected the nerves and allowed to retract |
| **Phantom position and mobility** | Phantom hand positioned slightly above the elbow; only feels the hand, not the forearm; can move all phantom fingers (Figure 1B). | Phantom hand positioned upright towards chest; only feels the hand, not the forearm; can move all phantom fingers (Figure 1B). | Phantom hand positioned upright towards chest; mostly hand and fingers (little elbow); can move all phantom fingers (Figure 1B). |
| **When did phantom sensations occur** | Immediately after amputation | Immediately after amputation | Immediately after amputation |
| **Phantom limb sensation (PLS) intensity** (100 max)<br>(3m, 6m, 1.5/5yrs respectively) | 40, 60, 40 | 90, 100, 100 | 100, 90, NA |
| **PLS frequency**<br>(3m, 6m, 1.5/5yrs) | 3m: once a week; 6m: several times per month; 1.5yr: once or less per month | 3m: all the time; 6m: all the time; 5yrs: all the time | 3m: all the time; 6m: daily |
| **Chronic PLS** (100 max)<br>(3m, 6m, 1.5/5yrs) | 13.3, 15, 8 | 90, 100, 100 | 100, 45, NA |
| **Limb pain intensity**<br>(Pre, 3m, 6m, 1.5/5yrs) | 90, 20, 0, 0 | 80, 50, 70, 70 | 50, 80, 70, NA |
| **Limb pain frequency**<br>(Pre, 3m, 6m, 1.5/5yrs) | Pre: all the time; 3m: several times per month; 6m: once or less per month; 1.5yr: once or less per month | Pre: all the time; 3m: daily; 6m: daily; 5yrs: all the time | Pre: daily; 3m: daily; 6m: once a week |
| **Chronic limb pain**<br>(Pre, 3m, 6m, 1.5/5yrs) | 90, 5, 0, 0 | 80, 25, 35, 70 | 25, 40, 23.3, NA |
| **Transient** (on the day) **limb pain**<br>(Pre, 3m, 6m, 1.5/5yrs; 100 max)<br>(Pre, 3m, 6m, 1.5/5yrs) | 50, 30, 0, 0 | 80, 45, 50, 70 | 50, 40, 20, NA |
| **Pain Detect Score**<br>(% max possible score)<br>(Pre, 3m, 6m, 1.5/5yrs) | 51%, 34%, 14%, 40% | 68%, NA, 42%, 45% | 65%, 65%, 65%, NA |
| **Pain Detect Pain Course** | - Persistent pain with pain attacks (Same pre and 3m)<br>- Persistent pain with slight fluctuations (6m, 1.5yrs) | - Persistent pain with pain attacks (Same pre and 6m)<br>- Persistent pain with slight fluctuations (5yrs) | - Pain attacks with pain between them (pre)<br>- Persistent pain with pain attacks (3m)<br>- Pain attacks without pain between them (6m) |
| **Upper Extremity Functional Index**<br>(Pre, 3m, 6m, 1.5/5yrs)<br>100% = no impairment | 47%, 23%, 36%, 57% | 30%, NA, 11%, 28% | 0%, 39% 69%, NA |
| **Prosthesis Type** | None | None (fitted with a cosmetic prosthetic) | Cosmetic prosthesis |
| **Prosthesis Use** | None | None. Briefly used in the first 6 months post-amputation (2 days a week, ~2 hours a day) | 6m: 2 days a week, 8 hours a day |

**Supplementary Table 2.** Pearson correlations for controls finger representations across sessions.

| Session Comparison | Hemisphere/Hand | Finger | Correlation Coefficient (r)<br>(mean ± std) |
| --- | --- | --- | --- |
| Pre to 3m | R/L | D1 | 0.83 ± 0.08 |
| Pre to 3m | R/L | D2 | 0.84 ± 0.10 |
| Pre to 3m | R/L | D3 | 0.88 ± 0.06 |
| Pre to 3m | R/L | D4 | 0.90 ± 0.04 |
| Pre to 3m | R/L | D5 | 0.89 ± 0.04 |
| Pre to 6m | R/L | D1 | 0.85 ± 0.07 |
| Pre to 6m | R/L | D2 | 0.84 ± 0.07 |
| Pre to 6m | R/L | D3 | 0.89 ± 0.05 |
| Pre to 6m | R/L | D4 | 0.90 ± 0.04 |
| Pre to 6m | R/L | D5 | 0.89 ± 0.04 |
| Pre to 3m | L/R | D1 | 0.80 ± 0.12 |
| Pre to 3m | L/R | D2 | 0.79 ± 0.13 |
| Pre to 3m | L/R | D3 | 0.83 ± 0.12 |
| Pre to 3m | L/R | D4 | 0.87 ± 0.07 |
| Pre to 3m | L/R | D5 | 0.88 ± 0.07 |
| Pre to 6m | L/R | D1 | 0.78 ± 0.16 |
| Pre to 6m | L/R | D2 | 0.79 ± 0.13 |
| Pre to 6m | L/R | D3 | 0.84 ± 0.10 |
| Pre to 6m | L/R | D4 | 0.87 ± 0.08 |
| Pre to 6m | L/R | D5 | 0.87 ± 0.07 |

## Methods

Our key methodology involves longitudinal comparisons across amputation. This approach is designed to overcome known limitations in cross-sectional designs, where inter-participant variability could spuriously influence group comparisons, particularly when considering small group sample sizes and/or small effects. An important additional consideration with respect to reorganization research in amputees is the difficulty to interpret whether sensorimotor activity for the missing (phantom) hand reflects preserved representation (i.e. reflects the same representational attributes as the physically hand prior to amputation), or an altered hand representation, which exhibits canonical hand representation features, albeit distinct from the pre-amputation hand. The main limitation of longitudinal designs is the contribution of any time-related effects, e.g. due to changes in MR scanning hardware^33^ or participants’ experience (e.g. familiarity with the study environment^34^, which are not directly related to the amputation. To account for non-related variables, we also scanned our case-studies and control participants over a similar timeframe. For two of our case-studies, we had an opportunity to follow up on our procedures after an extended period (1.5/5 years following amputation). As this was not planned in the original design, we were unable to obtain related timepoints in our controls. Therefore, all comparisons to the control cohort are focused on the 6 months point-amputation timepoint.

## Participants

### Longitudinal case-study participants that underwent an amputation

Over a 7-year period and across multiple NHS sites in the UK, we recruited 18 potential participants preparing to undergo hand amputations. Due to a multitude of factors (e.g., MRI safety contraindications, no hand motor control, age outside ethics range, high level of disability), we could only perform pre-amputation testing on 6 volunteers. Due to additional factors (complications during surgery, general health, retractions) we successfully completed our full testing procedure on 3 participants (for participant demographics see Supp. Table 1).

Pre-amputation scans for P1 and P2 were collected 24 hours apart and within 2 weeks of their amputations. P3 had a 2.5-year gap between the pre-amputation scans, due to Covid-related delays in testing and in scheduling uncertainty relating to their amputation surgery. Their amputation surgery took place 3 months following their second pre-amputation scan.

### Case-study participant amputation surgeries

There are noteworthy differences in their amputation surgeries of the three case-study participants. P1 underwent an amputation to combat a rapidly developing arteriovenous malformation (AVM) in the upper arm. Before amputation, they had a relatively high level of motor control in the pre-amputated hand. Additionally, P1’s amputation included more advanced surgical techniques, involving a combination of targeted muscle reinnervation [TMR]^35^ and regenerative peripheral nerve interfaces [RPNI]^36^. In these approaches, rather than simply cutting the residual nerve, the remaining nerves were sutured to a new muscle (TMR) or implanted with a nerve graft near a new muscle target (RPNI; in P1’s case, the technique varied depending on the muscle, see Supp. Figure 9). P2 underwent a traditional amputation procedure to remove a sarcoma tumor that had been slowly progressing since 1995. The multiple operations of the arm, prior to the amputation, left her with restricted motor control of the fingers, though still able to move them (see Supp. Video 1). Similarly, P3 was diagnosed with Severell-Martorell syndrome which had led to her left arm having multiple chronic bone fractures. They underwent a traditional amputation procedure, where the major nerves were left to naturally retract. It is important to note here that the diversity of conditions, procedures and post-operative states across our case-studies strengthen the universality of our results, which were consistent across case-studies.

### Longitudinal able-bodied control group

In addition to the case-study participants that underwent an amputation, we tested a control group which included 16 older able-bodied participants [9 females; mean age ± std = 53.1 ± 6.37; all right-handed]. The control group also completed four fMRI sessions at the same timescale as the participants that underwent an amputation and were age-matched to P2 and P3. 4 additional controls were also recruited for this group; however, we did not complete their testing, due to drop-out and incidental findings captured in the MRI sessions.

Ethical approval for all longitudinal study participants was granted by the NHS National Research Ethics Committee (18/LO/0474), and in accordance with the Declaration of Helsinki. Written informed consent was obtained from all participants prior to the study for their participation, data storage and dissemination.

### Cross-sectional datasets

From two previous studies^37^, we pooled two cross-sectional fMRI datasets: (1) a group of chronic amputees (n=26) and (2) a secondary group of able-bodied controls (n=18). The chronic amputee group included 26 upper-limb amputee participants [4 females; mean age ± std = 51.1 ± 10.6; 13 missing left upper-limb; level of amputation: 17 transradial, 8 transhumeral and 1 at wrist; mean years since amputation ± std = 23.5 ± 13.5]. The secondary able-bodied control group included 18 able-bodied participants [7 females; mean age ± std=43.1 ± 14.62; 11 right-handed]. For more information on these datasets, see Supplementary Methods (https://osf.io/s9hc2/).

### Longitudinal younger adults able-bodied control dataset

P1 is younger than the longitudinal control group. As such, we re-analyzed a previously collected dataset including 22 able-bodied controls of a similar age to P1 (mean ± std: 23.2 ± 3.8), each were scanned twice, one-week-apart on the same fMRI task and scanner^38^.

### Questionnaires

Due to a restricted time window for performing the tests before amputation, as well as the participants’ high level of physical discomfort and emotional distress, we were highly limited in the number of assessments we could perform. As such we focused the physically-involved testing on the functional neuroimaging tasks. However, in addition, we collected data on multiple questionnaires and had participants perform a functional ecological task.

#### Kinesthetic vividness

Kinesthetic vividness was quantified for each finger before and after the amputation [“*When moving this finger, how vivid does the movement feel? Please rate between 0 (I feel no finger movement) to 100 (I feel the finger movement as vividly as I can feel my other hand finger moving).”]*

#### Finger motor control

Perceived finger movement difficulty was quantified for each finger before and after amputation [“*When moving this finger, how difficult is it to perform the movement? Please rate between 100 (I found it as easy as moving the homologous finger in the unimpaired hand) to 0 (the most difficult thing imaginable).”]*.

#### Pain ratings

Before and after amputation, case-study participants were asked to rate the frequency of their pre-amputation limb pain or post-amputation phantom limb pain, respectively, as experienced within the last year, as well as the intensity of worst pain experienced during the last week (or in a typical week involving pain; see Supp. Table 1). Chronic pain was calculated by dividing worst pain intensity (scale 0–100: ranging from no pain to worst pain imaginable) by pain frequency (1 – all the time, 2 – daily, 3 – weekly, 4 – several times per month, and 5 – once or less per month). This approach reflects the chronic aspect of pain as it combines both frequency and intensity^39,40^. A similar measure was obtained for non-painful phantom sensation vividness and stump pain. Participants also filled out the Pain Detect questionnaire^41^. Additionally, before and after amputation, participants reported intensity values for different words describing different aspects of pain, quantified using an adapted version of the McGill Pain Questionnaire^42^. For each word, participants were asked to describe the intensity between 0 (non-existing) to 100 (excruciating pain) as it relates to each word.

Please note that we used a larger response scale than standard to allow the participants to articulate even small differences in their pain experience (see Supp. Figure 1).

#### Functional Index

Before and after amputation, case-study participants were asked to rate their difficulty at performing a diversity of functional activities because of their upper limb problem, quantified using the Upper Extremity Functional Index^43^.

### Ecological Task

To characterize habitual compensatory behavior, participants completed a task involving wrapping a present [based on ref. ^44^]. Task performance was video recorded but will not be reported in this paper.

### Finger Movement Task

To capture how participant’s move when cued to perform individual finger movements, at each session, we asked participants to perform a finger movement task where we cued them to move a single finger. Case study participants were cued to perform unilateral movements of the phantom fingers, intact fingers and then mirrored movements of the intact and phantom fingers simultaneously. Task performance was video recorded and is shown in Supp. Video 1.

### Intact Finger Kinematic Task

To test whether the intact fingers are being moved simultaneously during phantom finger movements, we invited 2 of the 3 case-study participants back for a separate session to assess the kinematics of the intact fingers. The task setup and data are shown in Supp. Figure 12.

### Scanning Procedures

Each MRI session for the longitudinal cohort consisted of a structural scan, four fMRI finger-mapping scans and two body localizer scans, which we report here. The additional cross-sectional datasets are detailed in the Supplementary Methods section.

### fMRI Task Design

#### Finger-mapping scans

The fMRI design was the same as a previous study from our lab^38^, though specific adaptations were made to account for the phantom experience of the case-study participants that underwent an amputation (described below).

Considering that S1 topography is similarly activated by both passive touch and active movement^24^, participants were instructed to perform visually cued movements of individual fingers, bilateral toe curling, lips pursing or resting (13 conditions total). The different movement conditions and rest (fixation) cue were presented in 9-second blocks and each repeated 4 times in each scan.

Additionally, each task started with 7 seconds of rest (fixation) and ended with 9 seconds of rest.

To simulate a phantom-like tactile experience for the participants pre-amputation, the affected hand was physically slightly elevated during scanning such that affected finger tapping-like movements were performed in the air. Alternatively, for the unaffected hand (before and after amputation), the individual finger movements were performed in the form of button presses on an MRI-compatible button box (four buttons per box) secured on the participant’s thigh. The movement of the thumb was performed by tapping it against the wall of the button box. For the control participants, half of the participants had the right hand elevated, performing the finger movements in the air, and the other half had the left hand elevated.

Instructions were delivered via a visual display projected into the scanner bore. Ten vertical bars, representing the fingers, flashed individually in green at a frequency of 1 Hz, instructing movements of a specific finger at that rate. Feet and lips movements were cued by flashing the words “Feet” or “Lips” at the same rate. Each condition was repeated four times within each run in a semi-counterbalanced order. Participants performed four scan runs of this task. One control participant was only able to complete 3 runs of the task for one of the sessions.

#### Imagery control scans

In each of the two body localizer scans, participants were visually cued to move each hand, imagine moving the affected (case-study participants) or non-dominant hand (controls), in addition to actual lips, toes (on the affected side only) and arm (on the affected side only) movements. The different movement conditions and a rest (fixation) cue were presented in 10-second blocks and repeated 4 times in each scan.

### MRI Data Acquisition

MRI images were obtained using a 3-Tesla Prisma scanner (Siemens, Erlangen, Germany) with a 32-channel head coil. Anatomical data were acquired using a T1-weighted magnetization prepared rapid acquisition gradient echo sequence (MPRAGE) with the parameters: TR = 2.53 s, TE = 3.34 ms, FOV = 256 mm, flip angle = 7°, and voxel size = 1 mm isotropic resolution. Functional data based on the blood oxygenation level-dependent signal were acquired using a multiband gradient echo-planar T2*-weighted pulse sequence ^45^ with the parameters: TR = 1.5 s, TE = 35 ms, flip-angle = 70°, multi-band acceleration factor = 4, FOV = 212 mm, matrix size of 106 x 106, and voxel size = 2 mm isotropic resolution. Seventy-two slices, with a slice thickness of 2 mm and no slice gap, were oriented parallel to the anterior commissure – posterior commissure, covering the whole cortex, with partial coverage of the cerebellum. Each of the four functional runs comprising the main task consisted of 335 volumes (8 min 22 s). Additionally, there were 204 volumes for the two imagery control scans (5 min 10 s). For all functional scans, the first dummy volume of every run was saved and later used as a reference for co-registration.

### fMRI Analysis

Functional MRI data processing was carried out using FMRIB’s Expert Analysis Tool (FEAT; Version 6.0), part of FSL (FMRIB’s Software Library, www.fmrib.ox.ac.uk/fsl), in combination with custom bash, Python (version 3) and Matlab scripts [(R2019b, v9.7, The Mathworks Inc, Natick, MA; including an RSA toolbox^46,47^. Cortical surface reconstructions were produced using FreeSurfer [v. 7.1.1^48,49^] and Connectome Workbench (humanconnectome.org) software.

Decoding analyses were carried out using scikit-learn (v.1.2.2).

### fMRI Preprocessing

The following pre-statistical processing was applied: motion correction using MCFLIRT^50^, non-brain removal using BET^51^, spatial smoothing using a Gaussian kernel of FWHM 3mm for the functional task data, grand-mean intensity normalization of the entire 4D dataset by a single multiplicative factor, and high-pass temporal filtering (Gaussian-weighted least-squares straight line fitting, with σ = 90 s). Time-series statistical analysis was carried out using FILM with local autocorrelation correction^52^. The time series model included trial onsets convolved with a double γ HRF function; six motion parameters were added as confound regressors. Indicator functions were added to model out single volumes identified to have excessive motion (>.9 mm). A separate regressor was used for each high motion volume (deviating more than .9mm from the mean position).

For the finger mapping scans, the average number of outlier volumes for an individual scan, across all participants, was 1.5 volumes.

To ensure all longitudinal sessions (Pre1, Pre2, 3m, 6m, 1.5/5 years) were well aligned, for each participant, we calculated a structural mid-space between the structural images from each session, i.e., the average space in which the images are minimally reorientated^53^. The functional data for each individual scan run within a session were then registered to this structural mid-space using FLIRT^50,54^.

### Low Level Task-Based Analysis

We applied a general linear model (GLM) using FMRI Expert Analysis Tool (FEAT) to each functional run. For the primary task, the movement of each finger/body-part (10 fingers, lips and feet – total of 12 conditions) was modeled against rest (fixation). To capture finger selectivity, the activity for each finger was also modelled as a contrast against the sum of the activity of all other fingers of the same hand.

We performed the same GLM analysis on the 6 conditions of the imagery scans. To capture the selectivity for actual attempted phantom movements versus imagine phantom hand movements, the activity for attempted hand movement was also modelled as a contrast against imagined hand movement.

For each participant, parameter estimates of the each of the different conditions (versus rest) and GLM residuals of all voxels were extracted from each run’s first-level analysis. All analyses were performed with the functional data aligned to the structural mid-space.

### Regions of Interest

#### S1: Broadmann Area 3b

We were specifically interested in testing changes in topography within (and around) BA3b. First, the structural mid-space T1 image were used to reconstruct the pial and white-gray matter surfaces using FreeSurfer’s recon-all. Surface co-registration across hemispheres and participants was conducted using spherical alignment. Participant surfaces were nonlinearly fitted to a template surface, first in terms of the sulcal depth map and then in terms of the local curvature, resulting in an overlap of the fundus of the central sulcus across participants^55^.

#### S1 (BA3b) hand region of interest

The BA3b ROI was defined in the fsaverage template space using probabilistic cytotectonic maps^55^ by selecting all surface nodes with at least 25% probability of being part of the grey matter of BA3b^56^. Further, for the multivoxel pattern analyses, we restricted the BA3b ROI to just the area roughly representing the hand. This was done by isolating all surface nodes 2.5 cm proximal/distal of the anatomical hand knob^57^. An important consideration is that this ROI may not precisely reflect BA3b for each participant and may contain relevant activity from neighboring S1 areas, due to the nature of our data (3T fMRI, smoothing FWHM 3mm) and the probabilistic nature of the atlas. As such, we consider this as a definitive localizer of S1 and an indicative localizer of BA3b. The surface ROIs were then mapped to the participant’s volumetric high-resolution anatomy.

#### 49 segments of BA3b

To segment BA3b into 49 segments, we loaded the fsaverage cortical surface with the boundaries of the BA3b ROI, as defined by the Glasser atlas^58^. We rotated the map so that the central sulcus was perpendicular to the axis. We overlayed a box with 49 segments of equal height, on this ROI. By masking the box to the ROI, we constructed 49 segments of the BA3b ROI. Because this masking approach requires drawing boundary lines using the vertices on the cortical flat map, we could optimally only get 49 segments (maximum) without issues with the boundary drawing approach. These ROIs were then mapped onto the participant’s volumetric high-resolution anatomy and further to the participant’s cortical surfaces.

#### M1: Broadmann Area 4

The approach for defining the motor cortex region of interest was the same as described above, with the sole exception of selecting the BA4 region.

### Projecting Functional Activity onto the Cortical Surface

Using the cortical surfaces generated using recon-all, fMRI maps were projected to the surface using workbench command’s volume-to-surface-mapping function which included a ribbon constrained mapping method. The only exception is the cross-sectional datasets where we projected all maps onto a standard cortical surface, see Supplementary Methods.

### Univariate Activity (in the order the analyses are reported across figures)

#### Contrast maps for moving versus imagine moving the phantom

To visualize the contrast maps for attempted versus imagine phantom hand movements, estimates from the two imagery-control scan runs for the participant’s post (6m) session were averaged in a voxel wise manner using a fixed effects model with a cluster forming z-threshold of 3.1 and family-wise error corrected cluster significance threshold of *p* < 0.05. Maps were then projected onto each participant’s cortical surface. These contrast maps are visualized in Figure 1C with a minimum z-threshold in both directions of 3.1.

#### Contrast maps for the hand and lips

To visualize the contrast maps for the hand and lip movements, estimates from the four finger-mapping scan runs for each session were averaged in a voxel wise manner using a fixed effects model with a cluster forming z-threshold of 3.1 and family-wise error corrected cluster significance threshold of *p* < 0.05. Maps were then projected onto participant’s cortical surface. These contrast maps (hand in red and lips in blue) are visualized in Figure 1D with a minimum z-threshold of 33% the maximum participant-specific z-statistic.

For completion, the boundaries of the lip maps, for all participants that underwent an amputation across all sessions, are visualized in Figure 3D. All maps were minimally thresholded at Z > 4.5 to provide a complementary thresholding approach relative to Figure 1D.

#### Hand topography across 49 segments of BA3b

Using the 49 segments of BA3b (described above), we projected the neural activity for the hand (versus rest) for each hemisphere (contralateral to the hand being moved), session and participant. The average activity across all voxels within each segment was averaged to extract a single value per segment.

#### Center of gravity

To quantify changes in the hand, finger or lip topography, we computed the center of gravity (CoG) of activity (for a single body-part) across the 49 BA3b segments. To do this, we first computed the weighted activity (β_w_) across the segments. To do this each segment number was multiplied by the average activity in the segment.

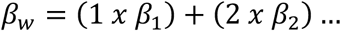

To compute the CoG, we then divided the sum of the weighted activity (∑β_w_) by the sum of the activity (∑β).

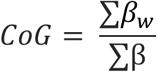

When comparing changes in the CoG for the hand or a finger, the CoG for each post-session was subtracted by the average CoG of the pre-sessions (e.g., 3m CoG – Pre. Avg CoG). A value greater than zero reflects the CoG moving more medially in the post session compared to the pre. A value less than zero reflects the post CoG being more lateral compared to the pre.

#### Finger selectivity maps

To visualize selectivity maps, estimates from the four finger-mapping scan runs for each session were averaged in a voxel wise manner using a fixed effects model. When visualizing the clusters, we minimally thresholded each z-statistic at 33% the maximum z-statistic. We stacked the images such that the smallest cluster is the highest overlay (e.g. the pinky) and the largest cluster is the underlay. Finally, we applied a 70% opacity to the visualizations to capture multi-finger activity at each voxel.

#### Representative control participant body-part maps

To provide an example visualization of the activity for each of the body-parts (shown in Figure 3C), estimates from the four finger-mapping scan runs for each session were averaged in a voxel wise manner using a fixed effects model with a cluster forming z-threshold of 3.1 and family-wise error corrected cluster significance threshold of *p* < 0.05. We then visualized the z-statistic map for the contrast of lips > feet and all left fingers > feet on an inflated cortical surface and applied a threshold to each body-part (Z > 3.1).

#### Lips activity in BA3b hand region

To test whether there is an increase in lip activity within the BA3b hand region, the average activity for all voxels (non-thresholded) in the ROI was computed for each session and each run. Activity was averaged across runs to compute a session estimate. When testing for a difference between the post and pre amputation sessions, the activity for the two pre-sessions was averaged for a pre avg. estimate. The activity in each post-amputation session (3m, 6m, 1.5/5y) was then subtracted to the activity of the pre avg.

### Winner-Takes-All Analysis

As a qualitative demonstration of our findings compatibility with previous studies investigating cortical reorganization that used a winner-takes-all approach, we applied a winner-takes-all analysis to S1 functional activity of the case-study participants that underwent an amputation. Using each participant’s final post-amputation session data, we performed two variations of the analysis including the conditions: (1) lips, hand and feet or (2) lips and feet (excluding hand). Each voxel was assigned exclusively to the condition with the highest activity. The resulting images were mapped to the participant’s cortical surface and visualized in Supp. Figure 10.

### Multivoxel Pattern Analyses

We performed several multi-voxel pattern analyses that can be broadly categorized into two themes: intra-finger, inter-finger and inter-body-part. In these measures, we were interested in capturing differences <u>within a session</u> and differences <u>between sessions</u>. For all of these analyses, we only included voxels within the BA3b hand region.

### Intra-finger

#### Pearson correlations

We first wanted to quantify changes in the pattern of activation for single fingers (intra-finger). We performed Pearson correlations on the beta-weights for each finger using data from runs from different sessions (Figure 2B; Supp Figure 5). For between-session correlations, the beta-weights [in our instance, contrast of parameter estimates (COPE)] for each finger in the 4 scan runs were separated into partitions each with 2 runs; each set from different sessions. The activity within each 2-run set were averaged at every voxel. A Pearson correlation was then performed between the averaged activity in each of the splits. We performed all unique 2-run combinations between-sessions (36 total combinations) and averaged these correlation coefficients to get a single value per finger. Between-session correlations were performed for all 6 unique session comparisons: Pre1 to Pre2, Pre1 to 3m, Pre1 to 6m, Pre2 to 3m, Pre2 to 6m, and 3m to 6m. Additionally, for P1 and P2, Pre1 to 1.5/5 years and Pre2 to 1.5/5 years. All correlation coefficients were then averaged and plotted in Supp. Figure 5. For a more simplistic visualization, we plotted just the first combination for each participant’s final scan relative to the Pre Avg. in Figure 2B.

#### Inter-finger

We next wanted to quantify changes in the pattern of activation between finger pairs (inter-finger) using a decoding approach (Figure 2D) and cross-validated Mahalanobis distances (Supp. Figure 6). Both approaches capture slightly different aspects of the representational structure^59^, which we elaborate on below.

For these two analyses, the beta-weights from the first-level GLM for each participant were extracted and spatially pre-whitened using a multivariate noise-normalization procedure [as described in ref. ^59^]. This was done using the residuals from the GLM, for each scan. We then used these noise-normalized beta-weights for the next analyses.

#### Decoding

First, we performed a decoding analysis. A strength of this approach is that it provides an estimate for chance performance (50%), i.e., *is the classification accuracy significantly greater than chance*. For the case-study participants that underwent an amputation, the decoding approach can tell us whether a decoder trained on pre-amputated finger pairs can correctly decode the same information on a phantom hand.

We used a linear support vector machine classifier (scikit-learn v.1.2.2; sklearn.svm, LinearSVC) to quantify between-session decoding for each finger pair. The default parameters were used for the classifier. Classification accuracy above chance (50%) denotes there is some amount of shared information between the train and test datasets.

We trained the classifier on the noise-normalized beta-weights for each finger pair (10 total). The train/test splits were performed using data from different sessions, such that the classifier was trained on each unique 2-run combination from one session and tested on all unique 2-run combinations in a separate session (36 combinations for each finger pair). We performed the same classification approach in the reverse direction (72 total combinations) because the forward and reverse directions provide unique values. The accuracies for each finger pair for each 2-run combination for each train/test direction were then averaged. Between-session accuracies are shown in Figure 1D.

#### Cross-validated Mahalanobis distances

Because our decoding analysis performed at ceiling (close to 100%), we also performed a representational similarity analysis using cross-validated Mahalanobis distances. The strength of this approach is that it computes a distance measure (continuous) as opposed to a binary decoding measure. As such, it is arguably more sensitive for capturing the inter-finger representational structure. Larger distances reflect more dissimilar (distinct) activity patterns and smaller distances reflect more similar patterns.

We performed this analysis using data from different sessions to compute between-session distances (our desired measure for representational stability over time). A distance cross-validated between sessions captures the stability of the information content.

We calculated the squared cross-validated Mahalanobis distance between activity patterns:

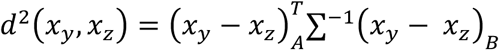

where (*x_y_* - *x_z_*) corresponds to the difference between the activity patterns of conditions y (e.g., thumb) and z (e.g., index finger) in partition A, and Σ refers to the voxel-wise noise covariance matrix. We performed this procedure over all possible 2-run cross-validation folds and then averaged the resulting distances across folds. There were 36 total unique cross-validation folds between-sessions.

We want to note that the cross-validated distance gives you the same distance value regardless of whether its assigned partition A or partition B. Between-session distances are shown in Supp. Figure 6.

#### Typicality

To quantify a measure that represents the degree of ‘normality’ of the hand representation, we computed a representational typicality measure^10^. For each participant’s non-dominant left hand, we extracted the 10 cross-nobis distances for the Pre-3m and Pre-6m comparisons. We then averaged these vectors across all the able-bodied participants to get an average typical hand pattern. We then performed a Spearman’s rho correlation between the cross-validated Mahalanobis finger-pair distances for each participant’s affected or non-dominant (left) hand and the average typical hand pattern. When comparing a control participant to the control mean, the respective participant was left out from the estimation of the control mean distances. These values are depicted in Supp. Figure 6.

#### Inter-body-part

Finally, we wanted to quantify changes in the pattern of activation between the thumb, lips and feet within the S1 hand region. We computed the cross-validated Mahalanobis distances between these body-parts in the same manner as the inter-finger analysis. The thumb to lips distances are plotted Figure 3. The distances between all conditions are plotted in Supp. Figure 7.

### Statistical Analyses

All statistical analyses were performed using either python scripts utilizing scipy.stats and statsmodels.stats.multitest or JASP (0.17.2.1). Tests for normality were conducted using a Shapiro–Wilk test. For the majority of analyses, to test whether a case-study participant was significantly different from the control group, we used Crawford and Howell’s method which provides a point estimate of the abnormality of each case’s distance from a control sample^60^. For all Crawford tests, we report uncorrected, two-tailed p-values. When comparing estimates to 0 or chance decoding (50%), we used a one-sample t-test (two-tailed). When testing for a decrease in measures within-participant, we used a Wilcoxon Signed-Ranks test. Additionally for the correlation analyses, Pearson correlations were used for the intra-finger multivoxel pattern analysis and Spearman correlations were used for the typicality analysis.

Across all of our previous studies, we operationally define amputees’ intact hand as their de-facto dominant hand, and as such have always compared non-dominant hand of controls to the missing hand of amputees (see for example refs.^9,14,40,61–64)^. Therefore, across all case-study to controls comparison analyses, we statistically compare (and plot) the controls left (non-dominant) hand side to the case-study participants missing hand side.

## Acknowledgements

We thank our participants for their immense generosity and dedication to contributing to this research. We thank the multiple clinicians that assisted in recruitment, namely: Dr. Imad Sedki, Dr. Stephen Kirker and Dr. David Henderson Slater. We thank Lina Teichmann, Hristo Dimitrov, Maryam Vaziri Pashkam and Raffaele Tucciarelli for feedback and support with analyses. We thank Clara Gallay for help with data collection.

## Author contributions

H.R.S. designed the research, collected the data, analyzed all datasets and wrote the manuscript. T.R.M. and C.I.B. designed the research, supervised analyses and edited the manuscript. M.K. helped collect data, preprocessed the cross-sectional datasets and edited the manuscript. M.A.S. helped collect data and edited the manuscript. R.O.M. designed the research, collected the data, supervised analyses and edited the manuscript. C.G., A.W., N.V.K. were involved in recruitment and editing the manuscript.

## Data and code sharing

Code and data used in the study will be made available following peer-reviewed publication.

